# NetMedGPT - A network medicine foundation model for extensive disease mechanism mining and drug repurposing

**DOI:** 10.64898/2026.01.04.697552

**Authors:** Farzaneh Firoozbakht, Simon Süwer, Maria Louise Elkjaer, Diane E. Handy, Andreas Maier, Jane Li, Lee Lancashire, Joseph Loscalzo, Jan Baumbach

## Abstract

Network medicine leverages large biomedical knowledge graphs (KGs) to model disease mechanisms and identify therapeutic opportunities. However, most deep learning approaches that use KGs in biomedicine remain task-specific, limiting their ability to generalize across diverse applications within a unified framework. Here, we introduce NetMedGPT, a transformer-based foundation model trained on a large-scale biomedical KG using masked token prediction. By learning contextualized representations of biomedical nodes, NetMedGPT enables unified, zero-shot inference across different drug discovery tasks. Specifically, in five tasks, i.e., predicting the association of drugs with indications, targets, adverse drug reactions, contraindications, and off-label uses, NetMedGPT consistently outperforms all specialized baselines, achieving area under the precision-recall curve gains of between 2.2% and 26%. When evaluated on independent external datasets, NetMedGPT outperformed baseline on an expert-curated clinical indications set and also preferentially prioritized clinically relevant drug-disease pairs in ClinicalTrials.gov. NetMedGPT’s generative capability further supports the construction of mechanistically plausible subnetworks offering biological insights. NetMedGPT provides a unified foundation model for network medicine that supports scalable hypothesis generation and provides potential to accelerate drug repurposing. We further provided an interactive interface (https://prototypes.cosy.bio/chatnetmedgpt/) that allows users to obtain model inferences through natural-language queries.

## Introduction

Network medicine views complex diseases as the result of perturbations across interconnected biological networks, requiring models that integrate molecular, phenotypic, and clinical information into a unified framework. Rather than attributing diseases to isolated genes or pathways, it emphasizes their origin in system-level disruptions within interconnected biological networks^1,2^. Therefore, therapeutic strategies should target the network context of disease rather than one or a few potential targets.

Network medicine seeks to elucidate disease mechanisms and to understand how therapeutic interventions perturb these networks and lead to clinical consequences^3^. One of its main translational applications is drug repurposing, which leverages network-based insights to uncover new therapeutic opportunities for existing drugs^1^. Such opportunities arise from the pleiotropic effects (off-target) of many drugs, which modulate multiple pathways beyond those involving their originally approved targets. These off-target effects can be detrimental, manifesting as adverse drug reactions (ADRs), or beneficial, revealing unexpected therapeutic potential. Repurposing harnesses the latter to reduce development time from approximately 13-15 years to around 6.5 years, and cost from around US $2-3 billion to an average of $300 million^4^, thereby lowering risk while expanding treatment options. A well-known example is imatinib, originally developed as a BCR-ABL inhibitor for chronic myeloid leukemia, which was later found effective in gastrointestinal stromal tumours by targeting the tyrosine kinase KIT (CD117)^5^.

Computational approaches, particularly network-based methods, have become central to advancing network medicine applications such as drug repurposing and therapeutic discovery^6–9^. By modeling both the topological structure (i.e., how nodes are connected) and the semantic context (i.e., what those connections mean), these models enable predictive reasoning across complex biomedical systems. Specifically, network-based approaches have shown promise in identifying novel therapeutic links. For example, baricitinib, an anti-inflammatory JAK inhibitor originally developed for rheumatoid arthritis, was identified via AI-based network analysis as a potential treatment for COVID-19 in early 2020. The prediction was subsequently validated in clinical trials, leading to FDA emergency use authorization in 2020 and full approval in 2022^10,11^.

In particular, for drug repurposing, many computational approaches are developed with the premise that effective interventions target biological networks perturbed by disease, either directly or through mechanistically relevant pathways^12,13^. Leveraging a drug-gene-disease knowledge graph (KG), Sadegh et al.,^14^ introduced a network-based drug repurposing framework (NeDRex) that identifies disease modules applying a range of module detection algorithms to seed genes, i.e., disease-associated or differentially expressed genes. Candidate drugs are then prioritized based on the network proximity of their targets to these inferred disease modules. To predict drug-disease links, Bang et al.^15^ introduced DREAMwalk, which uses a random walk guided by semantic similarity from drug and disease ontologies, enabling exploration beyond local graph structure. The resulting walk sequences are used to train a heterogeneous Skip-gram model to produce embeddings, which are then used as input to an XGBoost classifier that ranks drug-disease pairs. However, restricting KGs to only drugs, diseases, and genes overlook many important biological and clinical factors that influence therapeutic decisions. Recent efforts have addressed this gap by incorporating a broader range of biomedical nodes, such as ADRs, molecular functions, anatomical sites, and drug contraindications, into more comprehensive KGs. This expansion enriches the mechanistic landscape captured by the graph and enables more clinically relevant predictions. To facilitate this process, Chandak et al.^16^ compiled a multimodal KG for network medicine, called PrimeKG, integrating 20 biomedical data sources. Notably, PrimeKG includes drug-disease links for indications, contraindications, and off-label uses, making it a valuable resource for building more holistic repurposing models. Leveraging this comprehensive KG, Huang et al.^17^ introduced a graph neural network (GNN)-based model, denoted TxGNN, to predict drug indications and contraindications. By designing inference pipelines for zero-shot scenarios, suitable when no prior treatments are known or mechanistic knowledge is limited, TxGNN demonstrated state-of-the-art performance against several baseline models.

Another challenge in the context of network medicine is the reliable identification of drug-target interactions (DTIs). While pharmaceutical companies traditionally rely on labor-intensive and costly wet-lab experiments, computational approaches have emerged as efficient alternatives. In particular, leveraging KGs offers a powerful approach for drug target discovery as it enables the integration of heterogeneous biomedical data and the identification and prediction of undiscovered interactions^18–23^.

Similarly, predicting ADRs represents an essential frontier in network medicine. As clinical trials are constrained in the number of patients, duration of illness, and diversity, many ADRs remain undetected until post-approval when heterogeneous patient responses can reveal rare or long-term toxicities. For example, rofecoxib (Vioxx), which was approved for arthritis pain in 1999, was withdrawn in 2004 from the market due to serious cardiovascular risks, including an increased incidence of myocardial infarction and stroke that became evident only after wider clinical use^24^. Similar post-marketing concerns have emerged with drugs like rosiglitazone^25^ and varenicline^26^. To address this limitation, recent studies employ network-based methods to combine pharmacological and clinical data such that they can uncover latent drug-phenotype associations, facilitating earlier detection of ADRs and ultimately supporting safer drug development^27^.

In practice, identifying effective treatments requires incorporating multiple interdependent factors, including a drug’s mechanism of action, potential adverse effects, contraindications, and both approved and off-label indications. To provide meaningful translational value, computational models must, therefore, integrate and reason across these interconnected facets of pharmacotherapy rather than addressing them in isolation. Nevertheless, existing approaches mainly remain narrowly tailored to individual tasks, such as drug repurposing, limiting their capacity to reflect the complexity of clinical decision-making or to support holistic therapeutic reasoning.

A crucial aspect of biomedical AI is to provide biological insights, enabling researchers to uncover plausible mechanisms that support model outputs. Existing approaches typically rely on known biological relationships^17^, yet their utility is limited by the sparsity and incompleteness of current biomedical KGs in which many clinically or mechanistically relevant associations remain unreported. This limitation is particularly pronounced in early-stage translational settings such as with rare diseases (defined as affecting fewer than 200,000 people in the United States;^28^), polypharmacy (commonly defined as concurrent use of five or more medications^29^), or the detection of rare and under-reported ADRs (with incidence rates below 0.1% in interventional clinical trials^30^), where existing knowledge is fragmented. In these contexts, effective interpretability requires models that can infer and propose novel biologically plausible associations rather than relying solely on what already exists in the KG.

In this study, we introduce NetMedGPT, a transformer-based foundation model trained on graph-derived sequences using masked token prediction. NetMedGPT learns contextualized representations of biomedical nodes and edges, enabling unified zero-shot inference across diverse translational tasks. Without requiring task-specific retraining, NetMedGPT surpasses state-of-the-art performance on several tasks, including the prediction of drug indication, DTI, drug-ADR, contraindications, and drug-off-label use. In addition, NetMedGPT offers biologically meaningful insights into disease and therapeutic mechanisms, highlighting the promise of foundation models to bridge computational inference and clinical translation. Unlike existing approaches constrained by known edges in the KG, NetMedGPT generates context-specific subnetworks, revealing functional and potentially unreported biomedical associations. This generative capability supports the formulation of novel hypotheses and can guide wet-lab validation by highlighting relevant genes, pathways, or phenotypes. Moreover, NetMedGPT offers a scalable and cost-effective strategy for prioritizing repurposing candidates, helping to de-risk early-stage hypotheses and optimize the allocation of clinical trial resources.

## Results

### Overview of NetMedGPT for biomedical inference

Advancing network medicine requires models capable of reasoning across multiple dimensions of biomedical knowledge, such as drug mechanisms, adverse effects, indications, and contraindications, without the need for task-specific retraining. NetMedGPT is a transformer-based foundation model trained on a comprehensive biomedical KG to enable unified inference across a broad range of tasks (Fig. 1a). The model takes as input a masked sequence of biomedical nodes and edges, referred to as a masked pseudo-sentence, constructed from paths through the structure of the KG. In this context, each biomedical node and each edge type is treated as a distinct token in the model’s vocabulary. Next, it outputs predicted probabilities of the masked tokens. This operational strategy enables the model to make predictions across diverse biomedical inference tasks independent of the node and edge types. For example, in the drug repurposing task, the model receives a sequence containing a disease node and the indication edge and prioritizes candidate drugs. As the underlying KG, we use PrimeKG^16^, a publicly available biomedical resource comprising 129,375 nodes across 10 node types (Supplementary TableL1), connected by over four million edges across 30 edge types (Supplementary TableL2). In addition to its relational structure, we integrate prior knowledge about nodes by incorporating curated features from multiple biomedical databases (Supplementary Table 3, Fig.L1b).

**Fig. 1.**
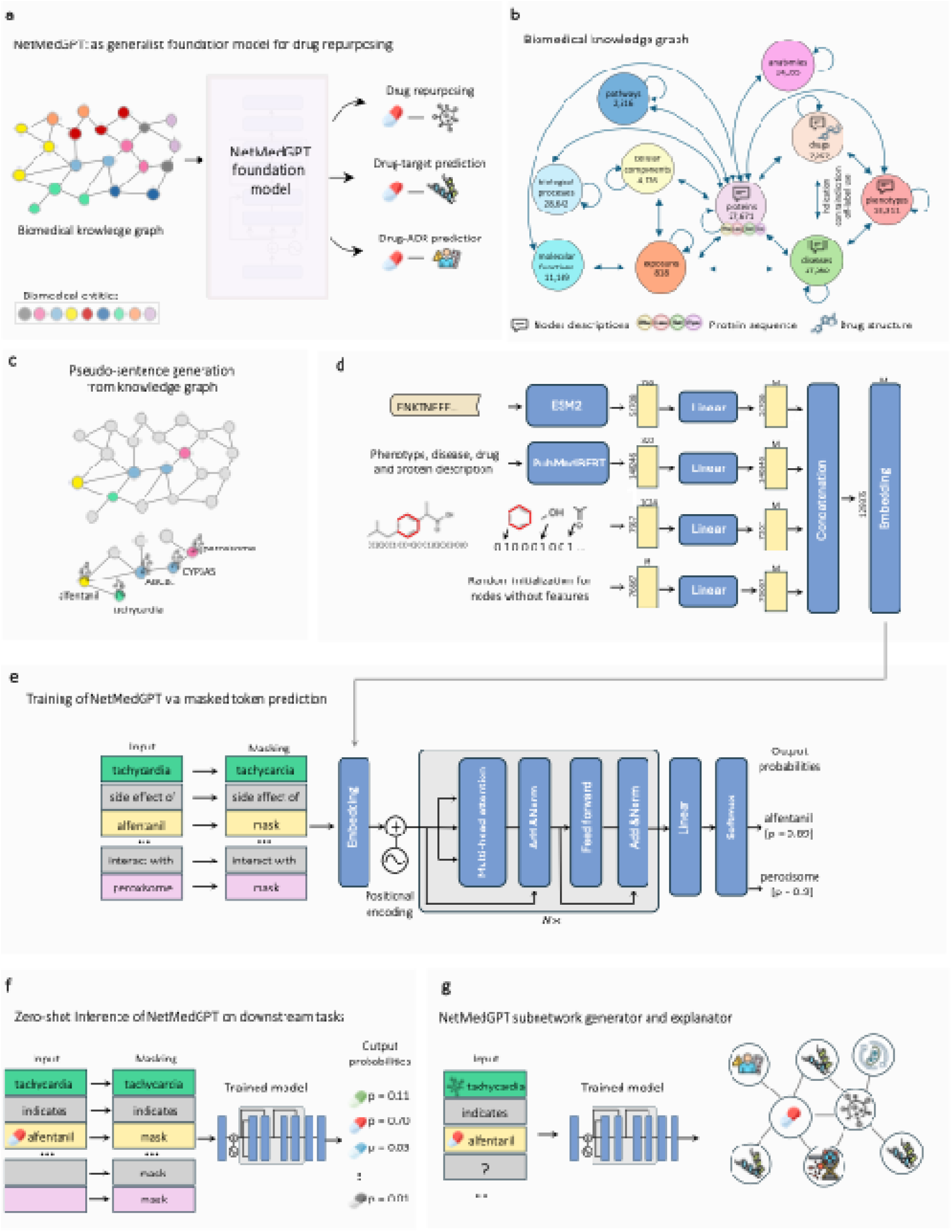
Overview of the NetMedGPT framework. a) NetMedGPT is a generalist, transformer-based foundation model developed to perform zero-shot inference across diverse drug discovery tasks, including the prediction of drug indications, protein targets, adverse drug reactions (ADRs), contraindications, and off-label uses. b) NetMedGPT is trained on PrimeKG, a large-scale biomedical knowledge graph (KG) encompassing 129,375 nodes and over four million edges spanning 10 node types and 30 relation types. Node-level features derived from curated biomedical databases are integrated for drugs, diseases, proteins, and phenotypes to enhance contextual representation. c) To train the model, pseudo-sentences are generated using random walks across the KG, capturing semantic meaning and network topology. d) NetMedGPT follows an encoder-only transformer architecture, including a token embedding layer, positional encodings, multiple self-attention layers, and a final linear layer for token prediction. During training, a subset of tokens in each pseudo-sentence is masked, and the model is optimized to predict the correct token using contextual information from the rest of the sequence. e) During inference, NetMedGPT is prompted with a task-specific pseudo-sentence containing masked positions and returns ranked predictions for the masked entity, enabling zero-shot predictions across tasks without further training. f) NetMedGPT generates subnetwork for each node pair by iterative next token prediction to provide mechanistic insight into biological processes.

The core principle underlying NetMedGPT is that biomedical nodes with similar neighborhoods in the KG tend to share functional relationships. For example, two drugs that share common target proteins and are associated with similar ADRs may also exhibit similar therapeutic effects. Capturing these implicit and often non-obvious similarities can be leveraged for different drug discovery tasks such as detecting new drug-disease associations.

To train NetMedGPT, we generated pseudo-sentences by performing random walks over the KG that capture both semantic and topological structure^31^ (Fig.L1c). Each pseudo-sentence is treated as a sequence of alternating node and edge tokens, capturing paths through the biomedical graph. Each token was associated with an initial feature vector. To combine node features, we incorporated drug molecular fingerprints derived from SMILES representations, pretrained embeddings from protein sequences (ESM-2), and pretrained textual embeddings from PubMedBERT for drugs, diseases, proteins, and phenotypes. For edge types and the remaining node types, we used learnable embeddings. All features were then aggregated to a shared embedding space (Fig. 1d). We then apply a masking pipeline that randomly hides a subset of tokens in each pseudo-sentence. A transformer-based model is trained to recover these masked tokens (Fig. 1e). The model uses self-attention to integrate information across all visible tokens in the sequence, enabling it to learn the statistical co-occurrence patterns, edge-type semantics, and higher-order topological dependencies among biomedical entities. In particular, by learning to predict a masked drug node, given surrounding disease, gene, and ADR context, the model implicitly learns multi-relational dependencies that support diverse downstream tasks. For example, in the pseudo-sentence of [*MASK, indication, type 2 diabetes, disease – protein, insulin receptor*], the model predicts different antidiabetic drugs as top candidates. At inference time, NetMedGPT is prompted with task-specific masked pseudo-sentences and predicts missing nodes or edges directly from the context, enabling zero-shot generalization across biomedical tasks without additional training (Fig 1f).

NetMedGPT adopts an encoder-only transformer architecture^32^, consisting of a token embedding layer, positional encoding, and a stack of *M* self-attention layers (Fig. 1e). Each input pseudo-sentence is tokenized into a sequence of integers, which are mapped into an initial embedding space. These embeddings can be initialized randomly or enhanced with prior biological knowledge. To incorporate the latter, we designed a modular embedding architecture that first extracts node features and then maps them into the same embedding space. Following the embedding and positional encoding layers, the model applies multiple self-attention layers to refine contextual representations, capturing higher-order dependencies and multi-relational semantics in the KG. Finally, a token prediction head, consisting of a linear followed by a softmax activation, is applied to predict probability distributions over the token vocabulary at each masked position. Additional architectural and training details are provided in Methods. By representing every node and edge as part of a shared vocabulary and training the model to predict masked tokens from their surrounding context, NetMedGPT learns general relational patterns that reflect the underlying structure of the KG. This design enables NetMedGPT to generalize beyond predefined tasks across a wide range of biomedical relations, distinguishing it from previous GNN-based models that are constrained to specific edge definitions.

Beyond predictive performance, it is essential for the model to provide mechanistic insight into biological processes. To this end, we leverage the generative capacity of NetMedGPT to identify subnetworks most contextually relevant to a given prompt query. Specifically, we developed a subnetwork generation framework based on next token prediction that iteratively expands an initial masked pseudo-sentence into a subnetwork (Fig.L 1g). For example, given a prompt consisting of a triplet (e.g. disease, indication, drug), we mask the remaining positions in a fixed-length sequence. NetMedGPT then performs iterative causal masked token prediction to fill in the missing tokens sequentially, generating a context-specific subnetwork of biomedical entities and relations inferred to be mechanistically relevant (Methods).

### Evaluation of NetMedGPT

We benchmarked NetMedGPT across a range of drug discovery tasks against state-of-the-art GNNs. In particular, as a baseline, we used relational graph convolutional networks (RGCN)^33^, heterogeneous attention networks (HAN)^34^, and a heterogeneous graph transformer (HGT)^35^. For drug indication and contraindication prediction tasks, we also included TxGNN. All models were evaluated based on a similar comparison framework on PrimeKG with consistent data split.

To evaluate model performance across different tasks, we used different cross-validation frameworks. Model performances are assessed using the area under the precision-recall curve (AUPRC), which quantifies the tradeoff between precision and recall and is well suited for imbalanced biomedical data as it emphasizes correct recovery of rare positives; the area under the ROC curve (AUC), which quantifies the tradeoff between true and false positive rates; and Recall@k, which quantifies the proportion of true positives within the top-k predictions.

### Generalist capability of NetMedGPT across network medicine applications

We evaluated the performance of NetMedGPT across multiple applications in network medicine, including the prediction of drug indications, ADRs, drug targets, drug contraindications, and off-label uses. We conducted an edge-level evaluation in which known links in the KG were randomly split into training, validation, and test sets (Fig. 2a). This setup allows us to evaluate the model’s ability to distinguish between positive edges, i.e., associations present in the KG, and negative edges, i.e., randomly sampled pairs with no known connection in the KG.

**Fig. 2.**
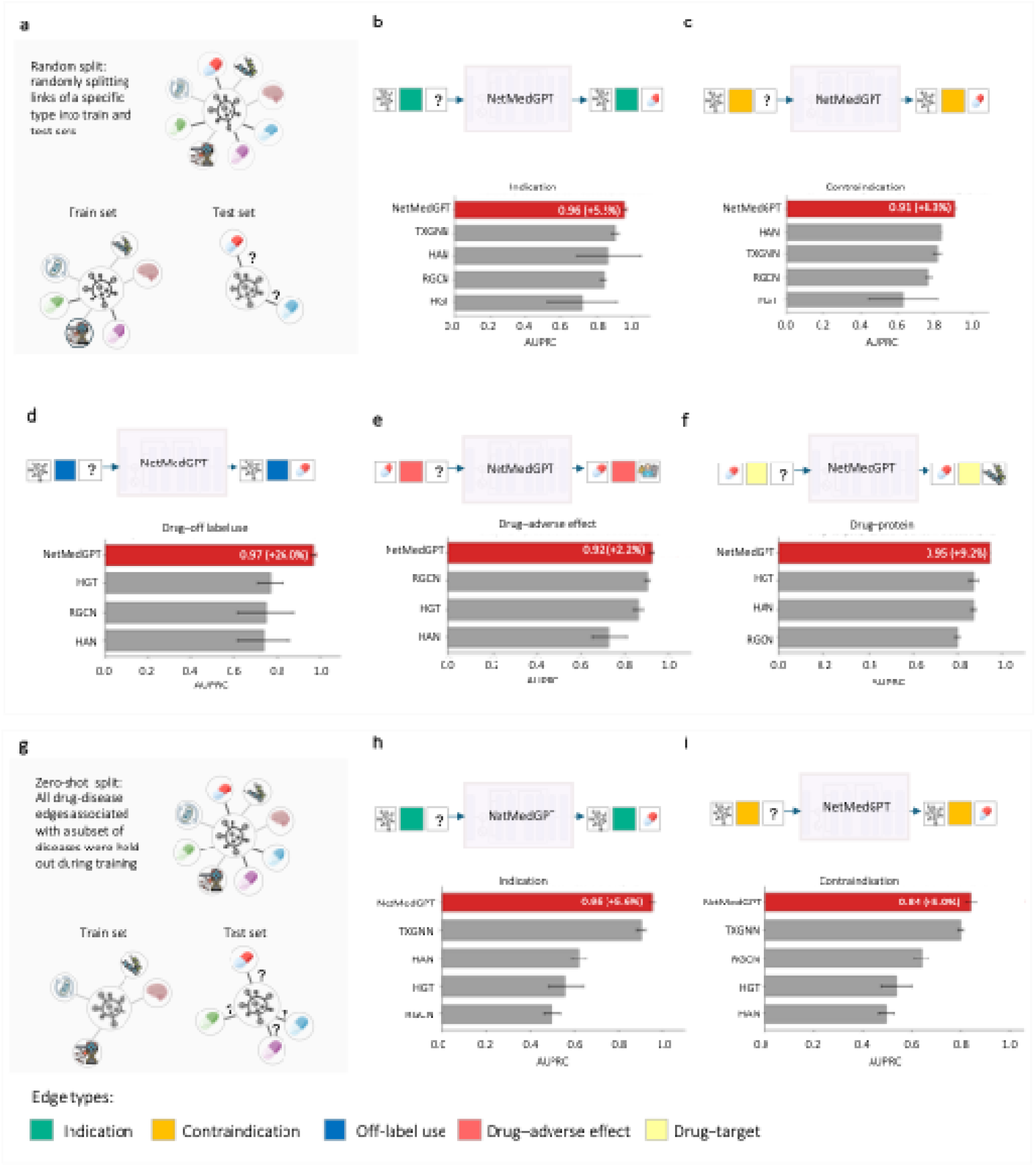
Evaluation framework and prediction performance of NetMedGPT across drug discovery tasks. a) Random link split evaluation: known edges in the knowledge graph (KG) are randomly divided into training, validation, and test sets. This setup evaluates the model’s ability to predict edges unseen during training. NetMedGPT outperforms the strongest baseline in b) drug indication prediction by 5.5%, c) drug contraindication prediction by 8.3%, d) off-label use prediction by 26.0%, e) adverse effect prediction by 2.2%, and f) drug-target prediction by 9.2%. g) Zero-shot split evaluation: all drug associations for a subset of diseases are held out during training, simulating real-world repurposing scenarios for diseases without approved treatments. Under this more stringent setting, NetMedGPT exceeds the best baseline in h) drug indication prediction by 5.6% and i) drug contraindication prediction by 5.0%. These results highlight NetMedGPT’s ability to infer clinically meaningful drug associations across both contexts, demonstrating strong generalization and reasoning in data-sparse settings. All AUPRC values are averaged over five independent runs (n = 5), with mean and standard deviation reported.

For drug indication, contraindication, and off-label prediction tasks, we adopted a disease-centric strategy^36,37^ to identify and prioritize drugs for a given disease. To this end, we prompted NetMedGPT with a pseudo-sentence of the form [*disease, indication, MASK*], where the model ranks candidate drugs at the masked position (Fig 2. b-d). Our results demonstrate that NetMedGPT significantly outperforms other baseline models across all three relation types. For indication prediction, TxGNN previously achieved state-of-the-art performance with an AUPRC of 91%, NetMedGPT improved upon this, reaching an AUPRC of 96%, improving performance by 5.5% (Fig. 2b). For contraindication prediction, the best-performing baseline was HAN, with an AUPRC of 84%. NetMedGPT surpassed this benchmark with an AUPRC of 91%, improving performance by 8.3% (Fig. 2c). In the off-label use prediction task, NetMedGPT achieved an AUPRC of 97%, outperforming the next best baseline, HGT, by 26.0% (Fig.L2d).

For ADR and DTI prediction tasks, the goal is to identify and prioritize relevant ADRs and protein targets for a given drug. To this end, we prompted NetMedGPT with pseudo-scentences of the form [*drug, indication, MASK*] and ranked candidate tokens at the masked position (Fig. 2e-f). As shown in Fig. 2e-f, NetMedGPT achieves an AUPRC ofL 92% for ADR prediction and 95% for DTI prediction, outperforming all baseline models by at least 2.2% and 9.2%, respectively.

While the improvements in AUPRC highlight NetMedGPT’s accuracy in ranking relevant entities among the top-scoring candidates, this metric is influenced by the choice of negative samples. In contrast, recall@k more directly reflects the model’s ability to recover true associations within the top-k-ranked predictions. As shown in Supplementary Fig.1, NetMedGPT achieves substantially higher recall@100 across all tasks, revealing its strong retrieval capacity compared to the baseline models.

To ensure a fair comparison with TxGNN, we adopted the same splits for the training, validation, and test sets, provided by Huang et al., ^17^. To further assess the robustness of our model, we also evaluated using other split configurations (Supplementary Fig. 2). As shown, the model’s performance (AUC and Recall@100) remained reasonably stable across these variations, exhibiting a gradual decrease as the proportion of training data was reduced.

We further assessed the contribution of node features to NetMedGPT’s performance through an ablation study. To this end, we re-trained NetMedGPT using only learnable node features without any prior knowledge-based features. As shown in Supplementary TableL4, removing those node features led to a consistent decrease in performance across all tasks, demonstrating that the incorporation of structured and text-based attributes enhances the model’s predictive power and generalizability. In addition, we conducted a stress test by systematically removing each edge type from the KG. This ablation led to moderate declines in performance demonstrating NetMedGPT’s robustness and its ability to generalize across incomplete relational contexts (Supplementary Fig.L3). Among all relation types, removing disease-disease links, derived from semantic similarities between MONDO ontology terms, had the most negative impact on model performance, particularly for indication, contraindication, and off-label use prediction. This underscores the critical role of semantic disease similarity in supporting mechanistic reasoning for drug discovery tasks. In contrast, removing drug-drug links, based on synergistic drug interactions, slightly improved performance. This finding suggests that such interactions contribute limited mechanistic information and may even introduce noise that impairs effective model training.

The baseline models are based on GNN architectures, which are known to struggle with capturing long-range dependencies due to issues such as over-smoothing, where node representations become indistinguishably similar across layers, and over-squashing, where rich information from distant nodes is compressed into limited-size embeddings^38^. In contrast, NetMedGPT addresses these limitations by leveraging graph-derived random walks and a transformer-based architecture. This design allows the model to attend to and integrate signals from distant nodes and across multiple relation types in a single forward pass, without being restricted to local neighborhood aggregation, thus enabling more expressive and global reasoning over the KG.

It is worth noting that, unlike NetMedGPT, which is trained in a unified task-agnostic manner, all baseline models were trained separately for each task. Our results show that NetMedGPT outperforms these task-specific models across multiple benchmarks. These results suggest the generalizability of NetMedGPT across diverse drug discovery tasks.

### Evaluating NetMedGPT in clinically motivated holdout scenarios

In the random link split scenario, the model’s capability is primarily evaluated on diseases that already have some approved drugs, aiming to identify additional candidates with potentially higher efficacy. To more rigorously assess model performance, we particularly consider a scenario where diseases do not have any known treatments. To this end, we applied the following two holdout strategies introduced by Huang et al.^17^.

i. *zero-shot split*: Predictions are made on diseases that were entirely unseen for that specific relation type during training. Specifically, all “indication” and “contraindication” edges for a randomly selected subset of diseases are withheld during training. The model must then predict treatments for diseases it has never seen linked to any therapy, although these diseases may still appear through other edge types.
ii. *disease-area split*: Predictions are made on diseases from a specific therapeutic area that are entirely held out during training. Specifically, all “indication” or “contraindication” edges corresponding to diseases within a specific biological category (disease-area) were held out for testing.

In both evaluation setups, the task of the model is to infer drug-disease associations for diseases that were not observed in the context of those specific edge types during training. For example, a disease may still be present in the KG via genetic or phenotypic edges, but all of its treatment-related edges (e.g., indications) are withheld for testing. The disease-area split imposes an even more stringent condition by removing all therapeutic associations for an entire disease category, requiring the model to infer plausible drug links based on shared biological mechanisms across disease classes, e.g., applying principles learned from metabolic diseases to predict drugs for cardiovascular conditions. Overall, these two setups evaluate the model’s ability to generalize beyond memorized associations, assessing whether it can reason over the observed biomedical contexts. Importantly, both setups can stress-test the model’s generalization capacity and its resistance to shortcut learning, where models rely on superficial statistical cues rather than meaningful biological patterns^39^.

In the zero-shot split setting, a subset of diseases is held out entirely for testing, while the remaining edges are divided into training and validation sets (Fig. 2g). As shown in Fig. 2h-i, NetMedGPT achieves AUPRCs of 95% for indication prediction and 84% for contraindication prediction, outperforming TxGNN, the strongest baseline, by 5.6% and 5.0% respectively. NetMedGPT also outperformed all other methods in terms of AUC and recall@100 (Supplementary Fig. 4).

To further evaluate the robustness of NetMedGPT, we conducted a series of stress tests. First, we randomly removed increasing fractions of edges from the KG (5%, 25%, 50%, 75%, and 90%) and monitored the resulting performance. While minor edge removals had minimal impact, performance declined significantly, by up to 26%, at higher removal ratios, as expected (Supplementary Fig. 5a,b). Next, we evaluated the model on a randomized graph, where NetMedGPT’s performance dropped to an AUPRC of 50%, indicating a random prediction (Supplementary Fig.L5c). Finally, to examine the impact of local biological context, we progressively masked 1-hop disease neighbors. Notably, NetMedGPT maintained robust performance even when 90% of the biological neighborhood was masked, achieving an AUPRC of 87%, demonstrating strong resilience to biological incompleteness (Supplementary Fig.L6).

For the disease-area split, we adopted the disease categorization from Huang et al.^17^ which groups diseases into nine clinically relevant categories: diabetes-related, adrenal gland, autoimmune, anemia, neurodegenerative, mental health, metabolic, cardiovascular, and cancerous disorders (Fig. 3a). The number of diseases varies across categories (Supplementary Table 5). For each category, all therapeutic edges associated with that category were excluded during training (Fig.L3b). The held-out category was then used exclusively for testing (Fig.L3c).

**Fig. 3.**
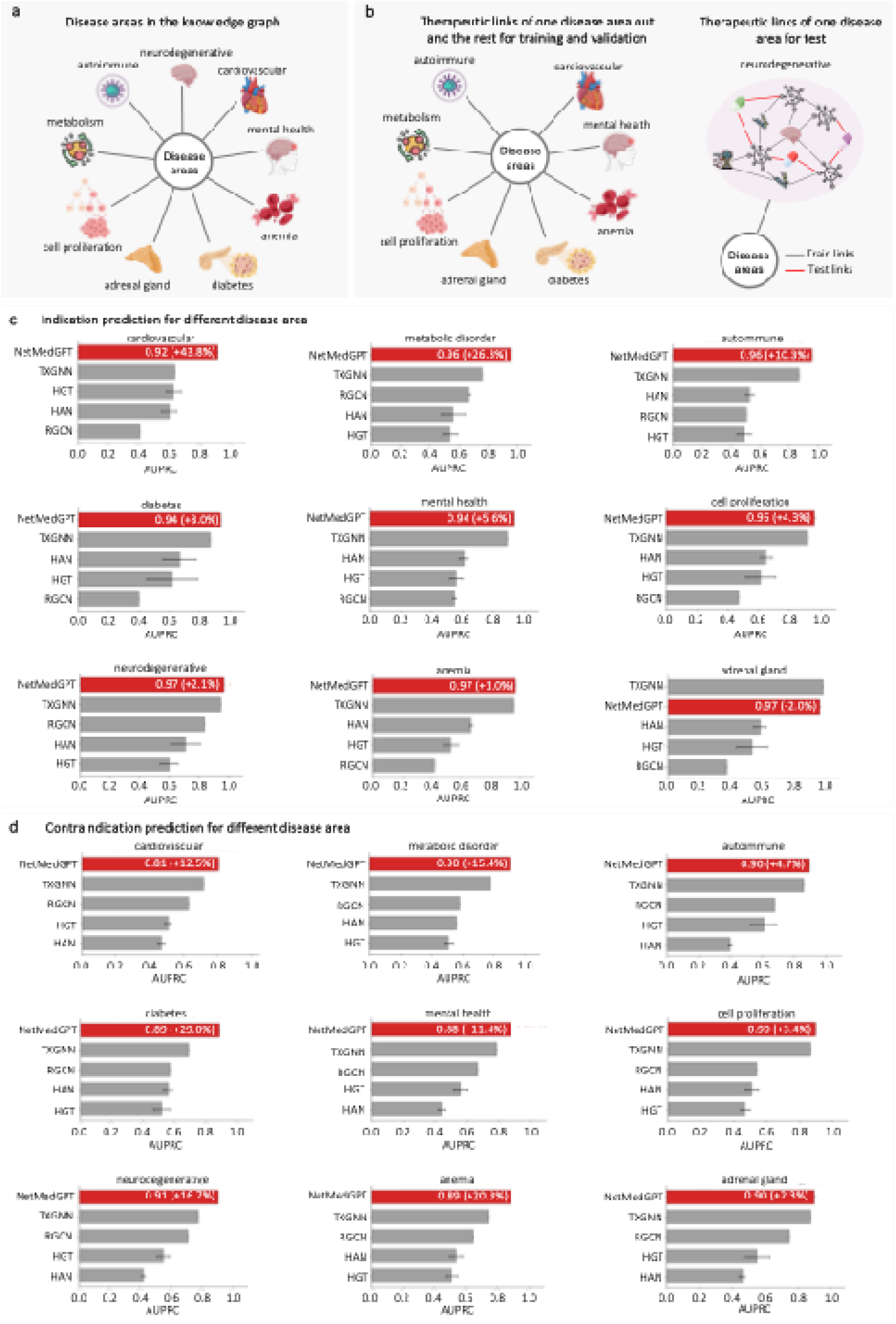
NetMedGPT performance on disease area-based indication and contraindication prediction. a) To evaluate generalization of the model, diseases were grouped into nine clinically relevant areas (following Huang et al. 2024): diabetes-related, adrenal gland, autoimmune, anemia, neurodegenerative, mental health, metabolic, cardiovascular, and cancerous disorders. b) For each area, all therapeutic edges were excluded from training and used as a test set. The remaining KG edges served for training and validation. c) For indication prediction, NetMedGPT outperformed all baselines in all disease areas except for adrenal gland disorders that was only 2% below the best model. d) For contraindication prediction, NetMedGPT achieved the best performance in all disease areas.

The comparison results for indication and contraindication prediction are shown in (Fig.L3c-d and Supplementary Fig. 7-8). For indication prediction, NetMedGPT achieved superior performance in 8 out of 9 areas (Fig.L3c) with an average relative AUPRC gain of 11% across all disease areas and a maximum gain of 43.8% in the cardiovascular area. In terms of recall@100, it outperformed other methods across all disease areas (Supplementary Fig. 7a). For contraindication prediction, NetMedGPT demonstrates even stronger generalization, outperforming all baselines in all areas, in terms of AUPRC (Fig.L3d), recall@100 (Supplementary Fig. 7b), and AUC (Supplementary Fig. 8b). It resulted in an average relative AUPRC gain of 12.8% and peak gains of 29.0% in diabetes.

### Evaluation of NetMedGPT using independent data

*Evaluation of NetMedGPT using a curated clinical indications dataset.* To increase confidence in our evaluation metrics, we further assessed the performance of NetMedGPT in comparison with TxGNN on an independent external dataset. We used a high-quality drug indication dataset obtained from Every Cure, a not-for-profit drug repurposing organization. This dataset is manually labelled by domain experts and includes both on-label and off-label indications, cross-referenced from multiple sources, including regulatory drug labels, clinical treatment guidelines, and the biomedical literature (Supplementary file 1).

The full dataset comprises 2,780 indication links covering 1,177 drugs and 795 diseases. We retained only indication links for which both drug and disease nodes were present in PrimeKG, resulting in 1,840 indications. We then excluded 716 indication links that overlapped with PrimeKG to ensure evaluation on previously unseen associations. The final evaluation set contained 1,124 indication links, spanning 583 drugs and 451 diseases, including 658 on-label and 466 off-label indications.

In our comparison analysis, we considered a TxGNN model previously trained based on a random link split setting, and trained NetMedGPT on the same training data. At inference, we generated ranked lists of candidate drugs for each disease in the curated dataset using both models and evaluated performance using recall@k (k = 1, 10, 20, 50, 100, 200) for on-label, off-label, and combined indications (Fig. 4a-d). Results show that NetMedGPT achieved significantly higher recall across all scenarios.

**Fig. 4.**
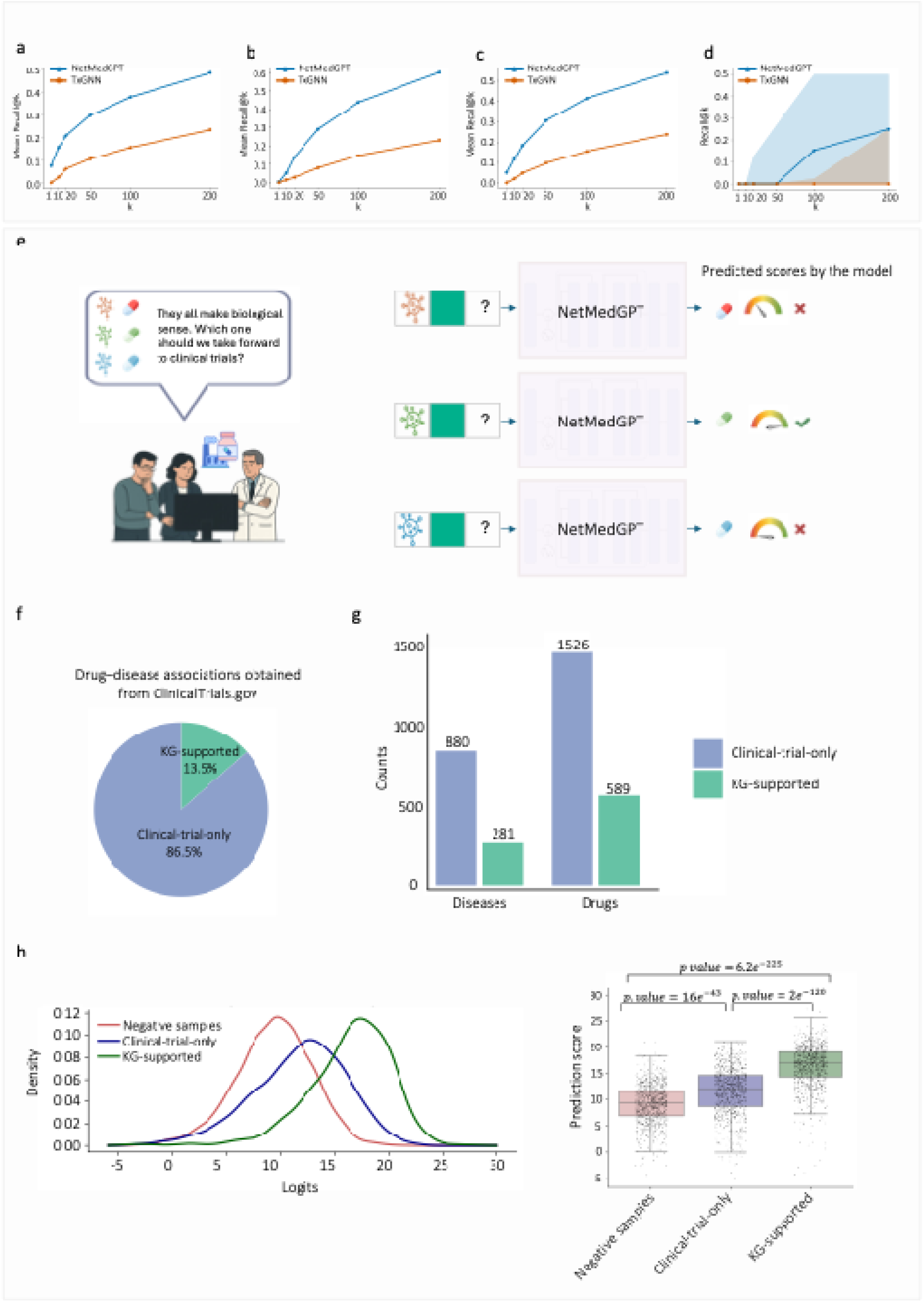
NetMedGPT in clinical decision support. Evaluation of NetMedGPT against TxGNN using an independently curated clinical indication dataset. For each disease in the curated set, all drugs were ranked and Recall@K was computed. Mean Recall@K is represented for a) on-label indications, b) off-label indications, and c) their combination. d) Median Recall@K is represented for the combination of on-label and off-labels indications; shaded regions show the interquartile range (Q1-Q3) across diseases, reflecting performance variability. NetMedGPT consistently achieves higher recall than TxGNN across all scenarios. e) NetMedGPT also provides a cost-effective strategy to prioritize repurposing candidates by assigning likelihood scores to drug-disease associations, helping researchers decide which candidates to advance to clinical trials. f) Drug-disease associations from ClinicalTrials.gov were stratified into two groups: KG-supported (green; 13.5%) and clinical-trial-only (blue; 86.5%). g) Counts of diseases and drugs within KG-supported and clinical-trial-only groups. h) Left: density distributions of NetMedGPT’s predicted logits for KG-supported (green), clinical-trial-only (blue), and random negative samples (red). Right: predicted scores for the same groups, showing that NetMedGPT significantly distinguishes KG-supported associations from clinical-trial-only and random negatives. This demonstrates NetMedGPT’s potential to highlight drug-disease associations with greater real-world therapeutic relevance.

*Evaluation of NetMedGPT using clinical trials dataset.* To assess the translational relevance of NetMedGPT and its potential utility in clinical decision support, we evaluated its predictions against external data from ClinicalTrials.gov^1^. In real-world settings, many drug-disease associations may appear biologically plausible, but only a subset can be prioritized for clinical evaluation due to cost, time, and regulatory constraints. In this context, NetMedGPT offers a scalable method to prioritize the most promising candidates for further investigation (Fig. 4e).

We curated a dataset of drug-disease pairs from ClinicalTrials.gov, encompassing FDA-approved therapies, off-label use, investigational use, and failed trials. In PrimeKG, drug-disease links labeled as “indication” reflect clinically supported use, including FDA-approved. Based on their overlap with PrimeKG, we classified ClinicalTrials.gov drug-disease associations into two groups: KG-supported, present in both PrimeKG and ClinicalTrials.gov, and clinical-trial-only, present only in ClinicalTrials.gov and often representing inconclusive or failed trials. The number of drug-disease associations in each group is shown in Fig.L4f, and the corresponding number of unique drugs and diseases is shown in Fig.L4g.

To avoid information leakage, we removed all overlapping KG-supported pairs from PrimeKG before training. As negative controls, we randomly sampled drug-disease pairs that were absent from both ClinicalTrials.gov and PrimeKG, using an edge-aware sampling strategy (Methods) and matched the number of positive pairs to ensure a balanced evaluation.

For each queried disease, NetMedGPT was prompted with pseudo-sentences of the form [*disease, indication, MASK*] and likelihood scores were generated for all candidate drugs at the masked position. As shown in Fig.L4h, both KG-supported and clinical-trial-only drug-disease pairs received significantly higher scores than negative controls, indicating that NetMedGPT effectively prioritizes clinically meaningful associations over random ones. Notably, KG-supported associations were assigned significantly higher scores than clinical-trial-only associations, likely reflecting the stronger evidence base encoded within the KG. Together, these results demonstrate NetMedGPT’s ability to rank both established and emerging therapeutic hypotheses, underscoring its potential as a cost-effective computational tool for early-stage drug discovery and repurposing before expensive experimental or clinical validation.

### Reducing noise in the KG

One of the major challenges in current KG is the high level of noise, including incomplete information and spurious links. Specifically, many drug-ADR links originate from inaccuracies in patient-reported data, leading to misleading associations. For example, patients who experience worsening of their underlying disease, whether related to drug efficacy or entirely unrelated to the treatment, may report the disease itself as an ADR. As some representative erroneous drug-ADR link appears in PrimeKG: Bosentan is incorrectly reported to have pulmonary arterial hypertension (PAH) as an ADR, even though it does not induce PAH under any circumstances. Minoxidil is incorrectly reported to cause angina pectoris, even though it does not directly induce angina; although an excessive drop in blood pressure can reduce coronary perfusion in patients with advanced coronary disease, this is not a direct adverse effect of the drug. Felodipine is incorrectly reported as contraindicated in coronary artery disease, even though it is clinically used to manage coronary artery disease (CAD)-related angina pectoris.

In addition, some phenotypes annotated as ADRs in the KG do not represent true drug-induced events. For example, in asthma, several reported ADRs, such as cough, wheezing, and nasal obstruction, are manifestations of the disease itself rather than effects attributable to pharmacological treatment. There are also cases in which drugs withdrawn from use for a specific indication remain linked to that disease in the KG. Voxelotor, a therapy designed to prevent sickle hemoglobin polymerization, was withdrawn by the FDA due to increased rates of vaso-occlusive crises and mortality, yet it still appears in the KG as a treatment for sickle cell disease. Conversely, important therapeutic links may be missing. For example, mavacamten, the first definitive therapy for hypertrophic cardiomyopathy targeting myosin mutants, is not included in PrimeKG.

Therefore, to alleviate noises in the KG, we performed a curation of the KG prior to generating subnetworks using insights from our analyses and established domain knowledge:

- **Removing drug-drug edges.** Our ablation analysis demonstrated that the existence of drug-drug edges decreases model performance in all tasks (Supplementary Fig. 3). This indicates that drug-drug edges in the KG may be affected by noise or bare redundant information; therefore, we removed these edges from the KG.
- **Filtering drugs with broad indications.** We observed that certain drugs, such as some glucocorticoids, are linked to a large number of indications. Since these drugs are often used to alleviate symptoms rather than to target specific mechanistic drivers, they are prescribed across many conditions, and their nonspecific usage artificially increases their connectivity in the KG and introduces bias into the learning process. Using an interquartile range (IQR)-based outlier detection, we identified and excluded these drugs from the KG (Supplementary Fig. 9).
- **Correcting misclassified drug-ADR edges**. Some drug-ADR edges arise because the drug is indicated for the same clinical term. Although rare cases exist in which a treatment can worsen the disease it is intended to manage (for example, via dose-dependent effects or toxicity), retaining all such links yields a large set of misleading ADRs driven by disease progression rather than true adverse effects. To mitigate this, we developed the following algorithm. First, we embedded all diseases and ADRs into a shared semantic space using PubMedBERT, which captures contextual similarity between biomedical terms based on large-scale literature pretraining. For each drug-disease pair with indication link, we computed the cosine similarity between the disease embedding and each ADR associated with the drug. Based on this semantic proximity, if any ADR associated with the drug appeared as the first nearest neighbor to the disease in the embedding space, we considered that drug-ADR link likely erroneous and, thus, removed it from the KG.

### Mechanistic insight through subnetwork generation

To provide mechanistic insight into its predictions, NetMedGPT introduces a generative reasoning framework that constructs biological subnetworks around predicted associations. Inspired by next-token prediction in generative language models such as GPT, the model starts from masked pseudo-sentences and iteratively completes masked positions by sampling from its learned probability distribution. For example, for drug-disease associations, the initial pseudo-sentences can be in the forms of [⌷⌷⌷⌷⌷⌷⌷⌷, *indication*, ⌷⌷⌷⌷] and [⌷⌷⌷⌷, indication, ⌷⌷⌷⌷⌷⌷⌷⌷]. The process of next token prediction is repeated multiple times to generate a set of contextually relevant biomedical nodes and edges to uncover plausible mechanistic subnetworks surrounding the initial association (Fig. 5). Accordingly, NetMedGPT moves beyond black box predictions by generating interpretable subnetworks that offer biological rationale for its outputs.

**Fig. 5.**
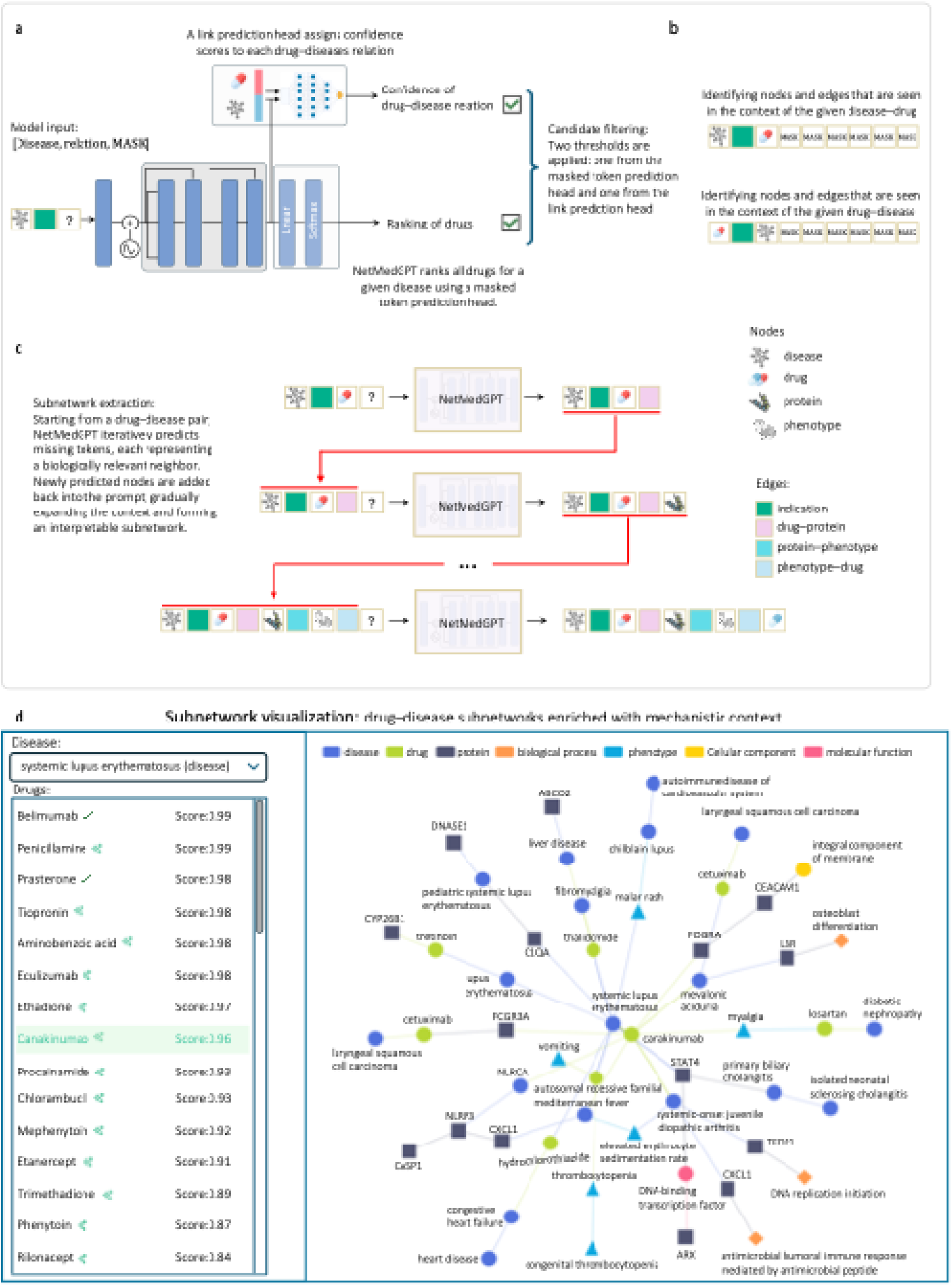
Subnetwork generation in NetMedGPT. a) NetMedGPT first identifies high confidence disease-drug associations by ranking nodes via the masked token prediction head and assigning confidence scores to each candidate pair by a link prediction head. b) The model then continues predicting MASK tokens to identify subnetworks around drugs and diseases. c) In the inference mode, the model starts from masked pseudo-sentences and iteratively predicts missing tokens to generate context-specific biomedical subnetworks. d) Our web interface enables users to interactively select diseases and prioritize drugs.

While masked token prediction enables ranking of nodes by applying softmax to assign relative likelihoods, it does not inherently produce calibrated probability estimates suitable for binary decision-making. To address this limitation, we added a link prediction head as the output of the frozen transformer encoder to assign calibrated probability scores to each pair (e.g., drug-disease) indicating how likely the model considers it a potential association (Fig. 5a; Methods).

We further retrained NetMedGPT on the preprocessed KG and applied next-token prediction to infer mechanistically relevant subnetworks around a given drug-disease pair. Due to the computational cost of subnetwork generation, we precomputed and provided subnetworks for ten representative diseases in the user interface (https://prototypes.cosy.bio/chatnetmedgpt/). The selected diseases were chosen to span diverse genetic architectures and therapeutic contexts, including complex polygenic disorders with multiple treatment strategies (asthma), endotype-driven diseases (coronary artery disease, COPD), monogenic conditions with multiple clinical phenotypes (sickle cell anemia, cystic fibrosis, hypertrophic cardiomyopathy), rare monogenic diseases lacking approved therapies (Angelman syndrome), and complex heterogeneous diseases characterized by immune, vascular, or neurodegenerative mechanisms (systemic lupus erythematosus, pulmonary arterial hypertension, Alzheimer’s disease).

Systemic lupus erythematosus (SLE) is a chronic autoimmune disease characterized by widespread immune dysregulation and multiorgan inflammation^40^. NetMedGPT identified mechanistically plausible candidates. IL-1 blockade (rilonacept, canakinumab) was linked to thrombocytopenia, and antiphospholipid antibodies, features in 5-50% of SLE patients^41–44^, alongside regulators such as STAT4, and HLA-DQA1, aligning with IL-1β-driven vascular inflammation and thrombosis^45^, and suggesting relevance in thromboinflammatory endotypes. These examples demonstrate NetMedGPT’s ability to recognize context-dependent immune regulation, enabling nuanced drug repurposing and endotype-specific mechanistic insight. However, the subnetworks should be viewed as mechanistic hypotheses rather than therapeutic evidence (Fig. 5d).

Coronary artery disease (CAD) is characterized by the narrowing or blockage of the coronary arteries most commonly due to atherosclerosis, resulting in reduced blood flow to the heart muscle. Statins, the standard therapy for lowering cholesterol, exert pleiotropic effects beyond their lipid-lowering action, which contribute to their overall therapeutic benefit, particularly when administered acutely during myocardial infarction. Notably, NetMedGPT retrieved a statin of lovastatin, as therapeutically relevant for CAD. In addition, clopidogrel, a P2YLL receptor inhibitor belonging to the class of antiplatelet agents, is commonly used in patients with CAD, particularly after stent placement, to prevent thrombosis, myocardial infarction, and stroke. Although the association between clopidogrel and CAD was not present in PrimeKG, it was recovered by NetMedGPT.

Multiple sclerosis (MS) is a chronic inflammatory disease of the central nervous system driven by immune-mediated demyelination and neurodegeneration^46^. When ranking candidate drugs for MS, NetMedGPT prioritized a large number of established disease-modifying therapies with high confidence score (>0.97) including both drugs annotated as indications in primeKG (e.g., teriflunomide, natalizumab) as well as several therapies inferred by the model despite missing indication links in the KG. These inferred high-confidence drugs included cladribine, alemtuzumab, fingolimod, orelizumab, and dimethyl fumarate, all of which are widely used in clinical practice^47^. The recovery of these therapies indicates that NetMedGPT captures key immunological and therapeutic features characteristic of MS and prioritizes clinically established treatments beyond KG annotations.

### Natural language-based NetMedGPT inference

To facilitate practical use of NetMedGPT, we implemented a user query-based inference pipeline that allows natural-language questions to be translated into structured model inputs (Fig. 6). Specifically, user queries are first processed by a large language model (LLM; GPT20b), which extracts biomedical entities and relations and maps them into a structured pseudo-sentence with one MASK token defining the prediction target. This step ensures that user queries are converted into a representation that is directly compatible with NetMedGPT’s input. To reduce the probability of misinterpretation, the structured pseudo-sentence is reconstructed into natural language and presented back to the user. Since user-provided entities, such as disease and drug names, may not exactly match node identifiers in the KG, we perform an explicit grounding step. To this end, user query entities together with all entities in the KG are embedded using PubMedBERT and the user query entities are mapped to their closest KG entities via a FAISS-based nearest-neighbor search^48^. Notably, this step enables robust alignment between user queries and biomedical entities represented in the KG, even in the presence of synonyms or lexical variation. Next, the grounded pseudo sentence is tokenized and fed to NetMedGPT to infer and prioritize candidate entities corresponding to the masked position.

**Fig. 6.**
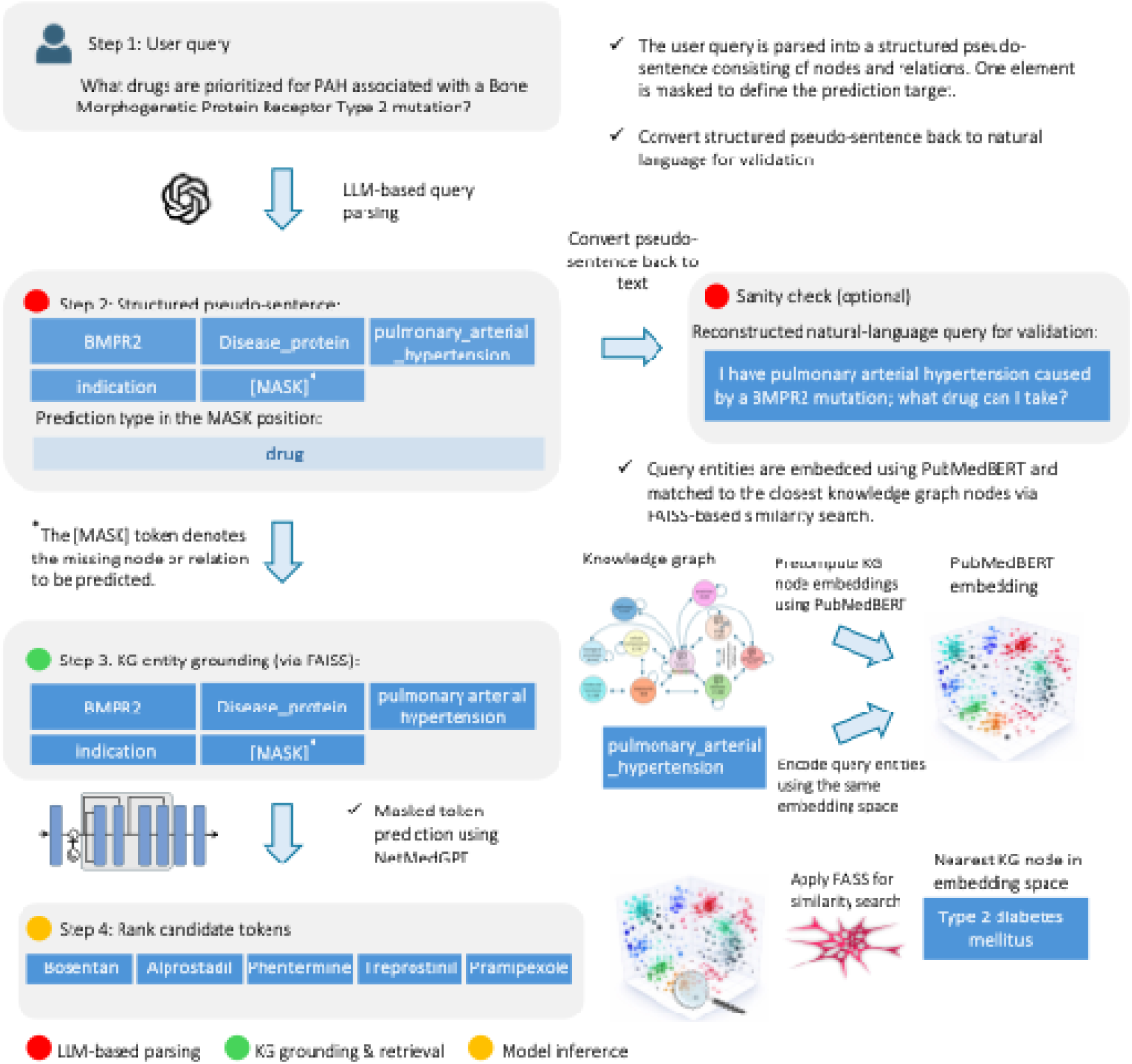
User query-based inference with NetMedGPT. User queries are first parsed by a large language model into a pseudo-sentence, compatible with the input of NetMedGPT, composed of nodes and relations with one element masked to define the prediction target. The pseudo-sentence can be optionally reconstructed into natural language to validate correct query interpretation. Query entities are then embedded using PubMedBERT and grounded to the closest knowledge graph nodes via FAISS-based similarity search in a shared embedding space. Next, NetMedGPT performs masked token prediction to infer candidate entities corresponding to the masked position.

## Discussion

A fundamental limitation of most computational methods developed for different network medicine applications is their narrow task specialization. Models are often developed and optimized for specific tasks, such as prediction of drug indications, DTIs, or ADRs, limiting their capacity for integrative reasoning. This siloed design becomes particularly problematic when the KG is too sparse for a given task. For example, in drug repurposing for rare diseases, where approved treatments are scarce and biomedical KGs are often incomplete, an effective approach requires integrating diverse and sparse biological evidence to generate robust therapeutic hypotheses.

NetMedGPT is a generalist foundation model designed to learn across diverse biomedical domains using a comprehensive KG. Rather than relying on task-specific supervision, NetMedGPT employs a transformer-based architecture trained with a masked token prediction objective to jointly learn all relations within the KG. Specifically, the model treats biomedical entities and their relationships as tokens in a shared vocabulary, allowing it to capture pharmacological, molecular, phenotypic, and clinical domains in a single framework. The model takes as input a pseudo-sentence composed of biomedical nodes and edge types as tokens, where a subset of tokens is randomly masked and must be predicted from context. Inspired by masked language modeling, this approach enables NetMedGPT to capture rich semantic dependencies across pharmacological, genetic, phenotypic, and clinical data. This capability is especially valuable for sparse or poorly annotated domains, including rare diseases.

We evaluated the performance of NetMedGPT based on different validation strategies. First, we applied an edge-level split, where a subset of edges of a specific type (e.g. disease-indication-drug) was randomly held out from the KG for testing. In this scenario, NetMedGPT consistently outperformed existing methods across different network medicine applications. To better simulate real-world drug repurposing, we further adopted a more stringent zero-shot split, in which all indication (or contraindication) edges for a subset of diseases were removed during training. This setup evaluates the model’s ability to identify potential treatments for diseases with no known treatment. However, partial information leakage may still occur if biologically related diseases appear in both training and test sets. To mitigate this, we implemented a disease-area holdout strategy where all drug-related edges (indications, contraindication, and off-label uses) associated with diseases from a specific clinical category were excluded from training. This setting provides a more rigorous benchmark that assesses the model’s capacity to generalize to entire disease groups with no prior treatment information, representing a more clinically meaningful repurposing evaluation. Notably, across all evaluation scenarios, NetMedGPT demonstrated not only superior mean performance but also small variance across replicates, which indicates its robustness and consistent generalization across diverse biomedical contexts.

To evaluate its real-world utility, we tested NetMedGPT’s predictions against external evidence from <u>ClinicalTrials.gov</u>. The model significantly differentiated KG-supported from clinical-trial-only and random drug-disease associations, indicating its ability to prioritize clinically meaningful relationships. Our results demonstrate the potential of NetMedGPT as a hypothesis generation tool for early-stage drug repurposing. However, we note that these predictions require expert curation and experimental validation and should not be interpreted as verified clinical advice.

Given the growing recognition that complex diseases often arise from disruptions across multiple pathways^49^, there is a need for models that can reason beyond single-target therapeutics. Building on NetMedGPT’s generative reasoning framework, future directions include identifying synergistic drug combinations that target distinct yet complementary pathways associated with a disease. This extension could enhance its utility in designing synergistic treatment regimens, especially in multifactorial or treatment-resistant conditions, aligning computational predictions more closely with emerging paradigms in network-based and systems pharmacology.

NetMedGPT’s performance is inherently dependent on the quality and coverage of the underlying KG, i.e., PrimeKG. Like all curated resources, PrimeKG may contain incomplete information, such as missing drugs, underrepresented disease areas, or biased edge distributions. However, sparsity is not merely a data limitation but also an intrinsic characteristic of biomedical KGs. Even with complete biological information, the space of known drug-disease associations would remain inherently sparse. In such settings, the model learns to identify alternative paths connecting drugs and diseases, often revealing mechanistically relevant candidates whose directionality may still require further investigation. Although we implemented a post-processing step for the refinement of drugs prioritized for a disease, future improvement is required to enhance the practical utility of NetMedGPT optimally.

In addition, our KG currently lacks patient-level data, such as omics profiles (e.g., transcriptomics, proteomics, and genomics), electronic health records (EHRs), demographics, and medical imaging. Incorporating these data modalities could allow more personalized predictions by accounting for individual variability, disease subtypes, and real-world clinical features. Future efforts to expand the KG could improve the accuracy, robustness, and breadth of NetMedGPT’s predictions.

In conclusion, NetMedGPT illustrates how foundation models can unify biomedical knowledge to improve therapeutic discovery. Its framework could be extended to support biomarker discovery and personalized medicine, thereby informing more efficient trial design. NetMedGPT represents an initial step toward generalist AI systems for biomedical reasoning.

## Methods

### Dataset

We used PrimeKG (Chandak, Huang, and Zitnik, 2023) as the foundational biomedical KG for training NetMedGPT. PrimeKG is a publicly available, multimodal, large-scale KG specifically curated for precision medicine applications. It integrates data from diverse biomedical resources and contains over 129 million nodes spanning 10 node types, along with more than 8 million edges across 30 edge types (Supplementary Table 1). For downstream evaluation, we additionally incorporated data from over 300,000 clinical studies extracted from the ClinicalTrials website (*clinicaltrials.gov*). These data were used to identify emerging or clinical-trial-only drug-disease associations that are not present in structured indication or contraindication databases. This allowed us to assess NetMedGPT’s ability to predict novel therapeutic hypotheses beyond those represented in curated KGs, reflecting real-world drug target identification and repurposing scenarios currently under clinical investigation.

### Task definition and learning objective

We define a heterogeneous biomedical KG, *G* = (*V,E,R,T*), where *V* is the set of nodes {*v_i_*} representing biomedical nodes, E is the set of edges {*e_i_*}, *e*; = (*v_j_*,*v_k_*), with *v_j_*, *v_k_*, ∈ *V*. *T* is the set of node types, and each node *v_i_* ∈ *V* is assigned a type *t_i_* ∈ *T*. *R* is the set of edge types and each edge is assigned a type *r_i_* ∈ *R*. We then define a pseudo-sentence of length *L* as a random walk over *G*, consisting of *N* nodes and *N* — 1 edge types (2*N* — 1 = *L*) in the ordered tuple of (*v_i_*, *r_n_*, *v_j_*, *r_m_*, … ). Each element, whether a node or an edge, in the pseudo-sentence is a token indicated by an integer number. We further define a special token MASK that can replace any token in the pseudo-sentence to simulate missing information. Given a pseudo-sentence including several MASK tokens (< *L*), the model is trained to recover the original tokens at the masked positions. These masked elements can be either nodes or edges and may appear at any position in the sequence. We note that the complete vocabulary, therefore, consists of I *V* I + *R* | +1 tokens: one for each node, one for each edge type, and one special MASK token.

### Pseudo-sentence generator

To prepare data for training and evaluation, we generated sequences of pseudo-sentences from our KG. To this end, we used an undirected random walk algorithm introduced in node2vec^31^, which captures both local and global graph structure by biasing the walk behavior through hyperparameters of *p* and *q*. Given a starting node *v*, a random walk is performed by iteratively transitioning to a neighboring node with a transition probability:

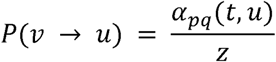

*z*, is a normalization constant ensuring the probabilities sum to 1, and α*_pq_*(*t*, *u*) is a bias factor that modulates the transition probability based on the structural properties of the graph. The bias factor is defined as:

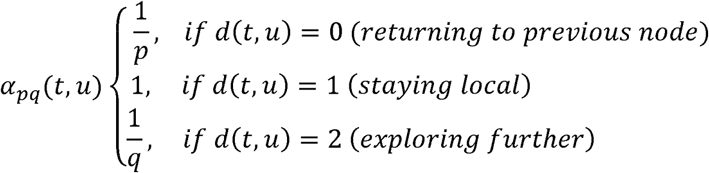

where *d*(*t, u*) denotes the shortest path distance between the previously visited node *t* and a candidate target node *u*. By adjusting *p*, the return parameter, and *q*, the in-out parameter, the algorithm balances between a breadth-first search, for capturing local neighborhoods, and a depth-first search, for exploring global graph connectivity that explore distant regions of the graph. In this work we considered *p* = 1 and *q* = 1 as the default settings used on the original implementations of node2vec^31^.

To ensure sufficient coverage of the graph structure, each node serves as the starting point for *N* independent random walks; each continues until a predefined walk length *N* is reached. The resulting sequences of nodes were then converted into pseudo-sentences by inserting the edge type between consecutive pairs of nodes in the sampled walks (Supplementary Fig 10). These sequences, representing heterogeneous multi-hop paths through the biomedical graph, serve as pseudo-sentences for the masked token prediction task, allowing NetMedGPT to learn contextualized embeddings of both nodes and edges. We set the walk length *N* = 5 and subsequently *L* = 9, indicating the context size of our model. The number of walks per node was set to 30.

### Model architecture

Each token in a pseudo-sentence, representing either a biomedical node or an edge type, is embedded into a learnable vector. These embeddings are then fed into a multi-layer transformer encoder that applies self-attention over the sequence to learn contextualized embeddings. The overall architecture of NetMedGPT is illustrated in Figure 1, which is composed of three parts described below.

#### Initial token embedding

Given a pseudo-sentence obtained by a random walker as input, each dimensional space. For certain node types (i.e. proteins, diseases, drugs, and phenotypes), we leveraged *a priori* knowledge by incorporating features derived from specialized embedding methods previously developed in the literature. These features were treated as fixed embeddings during training. The features and embedding sources used for each node type are listed in Supplementary Table 3. These features have various dimensionalities, which needs to be equalized as embedding vectors. To this end, we passed each feature set through a trainable linear layer, mapping them into the shared d dimensional embedding space. For node types of proteins and drugs, multiple features are used. Hence, for these nodes, separate linear layers were applied to each feature type, and the resulting vectors were summed to form the final embedding. For the rest of node types as token embeddings were randomly initialized and optimized during training.

#### Positional embedding

Following Vaswani et al.^32^, we incorporated sinusoidal positional embeddings to encode the order of tokens within each pseudo-sentence. For each token position *i* and embedding dimension index pos, the positional encoding is defined as:

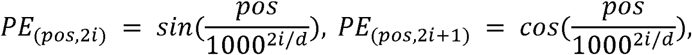

where *d* is the dimensionality of the model’s embeddings. These position-dependent vectors are added to the token embeddings to inject sequential information into the model, enabling it to distinguish token order within graph-derived sequences.

#### Transformer-based encoder

The embedding vectors generated by token embedding and positional embedding were summed and fed into K stacked transformer layers.

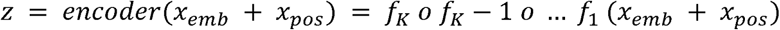

Each *f_i_* denotes a standard transformer layer composed of multi-head self-attention, feed-forward networks, layer normalization, and residual connections. We use the original transformer architecture introduced by Vaswani et al.^32^, implemented in PyTorch v2.2.0.

#### Token prediction unembedding

The encoder is followed by a decoder to perform mask token prediction. To this end, the embeddings generated by the encoder are fed to a linear layer with outputs followed by a softmax layer to convert logit values to probabilities. Formally, the output values are calculated as

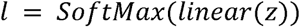

### Input masking strategy

During training, a subset of tokens in each pseudo-sentence is randomly masked and must be predicted by the model based on the given context. Given a pseudo-sentence, we applied masking with a probability of 0.2 per token, subject to the following constraints: (i) each sequence contained at least four masked tokens, even if fewer were selected by the random process; and (ii) at least one token was always left unmasked, preventing the model from seeing an entirely masked sequence. Masked tokens were then replaced with a special token identifier.

### Loss function

Training was performed using a cross-entropy loss computed over the masked tokens in each pseudo-sentence. The cross-entropy loss encourages the model to assign high probability to the correct token among all possible candidates in the vocabulary, thereby enabling it to learn contextualized representations of biomedical concepts. Let *S* = (*t*_1_, *t*_2_,…,*t_L_*) be a pseudo sentence of tokens sampled from the KG, where each token *t_i_* corresponds to either a biomedical node or an edge. A subset of positions M ⊂ {1, …, *L*} is selected based on masking strategy, described above, and replaced with a special [*MASK*] token. For each masked token, *t*; ∈ *S*, the model outputs a probability distribution *p_θ_* (*t_i_* | *S*/*M*) over the vocabulary. The objective is to minimize the negative log-likelihood of the correct token across all masked positions:

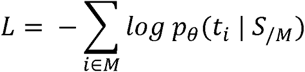

Where, *S*/*M* denotes the pseudo-sentence with the masked tokens, θ represents model parameters, *t_i_* is the true token at position *i*.

### Cross-validation and evaluation setup

To assess the model’s performance, we employed three cross-validation strategies:

*Edge-level cross-validation:* this validation scenario was applied to all drug discovery-related tasks, including the prediction of indication, contraindication, off-label use, drug-target and drug-ADR. Specifically, for each edge type, edges are randomly divided into train, validation and test with the ratio of 90-5-5, respectively.

*Zero-shot edge-specific node removal:* this validation scenario was applied only for the drug repurposing task that includes the edge types of indication and contraindication that link diseases to drugs. For each edge type, we first list diseases with at least one edge of that type. Then, we randomly divide these diseases into train, validation and test with the ratio of 90-5-5, respectively. Finally, all edges linked to diseases of each group were assigned to train, validation and test accordingly. This validation schema simulates zero-shot inference on unseen diseases for indication and contraindication edges.

*Disease-area generalization:* this validation scenario was also applied only to the edge types of indication and contraindication that link diseases to drugs. First, all diseases were grouped into categories of their area based on^17^. Then, for each disease area, all edges of a specific type were considered as a test and removed from the network. Finally, we used the zero-shot edge-specific node removal schema to divide remaining edges into train and validation.

### Negative sampling

To generate negative samples, we employed an edge-aware, degree-matched sampling strategy to represent realistic negative samples rather than implausible associations which is important for robust evaluation in biomedical link prediction tasks.

For each positive edge (head, tail) of type t in the evaluation set, we sampled an equal number of negative edges for the same head, subject to two constraints:

1. *Edge type-awareness*: the tail of negative edges was drawn only from nodes known to participate in the same edge type r elsewhere in the KG. For example, when sampling negative drugs, as tails, for a given disease, as the head, under the indication edge, we selected only drugs that are indicated for at least one other disease, ensuring the negative examples remain biologically relevant.
2. *Degree matching*: to preserve the degree distribution across node types, for each head node h, the number of sampled negative examples matched its number of positive links under the edge r.

### Subnetwork generation

In order to provide interpretability, we developed a framework that generates biologically plausible subnetworks conditioned on a given pseudo-sentence, which can simply be a triplet of *node_i_, edge_j_, node_k_*. For a pseudo-sentence of length *L*’ (here *L*’ = 3), we construct a full-length sequence of L tokens by masking the remaining *L* — *L*’ positions. The model then sequentially predicts high probability tokens for the *MASK* positions, iteratively completing the sentence until all masked tokens are filled (Figure 5c). This generation process is repeated R times to produce a diverse set of candidate subnetworks.

To select a high probability token at each step, we first extract the top-5 candidate tokens based on their logit scores. We then apply a temperature-controlled selection probabilities:

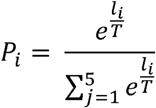

where *l_i_* is the logit score for the i,, token, and T is the temperature hyperparameter. Increasing *T* promotes greater diversity in the sampled tokens. We set *T* = 10 to allow exploration of multiple plausible biological completions while still focusing on high-confidence predictions.

### Link prediction head

To assign confidence scores to candidate drug-disease associations, NetMedGPT employs a link prediction head, a multi-layer feedforward neural network trained as a binary classifier. Given the contextual embeddings of the two target nodes (e.g., a drug and a disease), their representations are concatenated and passed through a series of fully connected layers with activations to predict the likelihood of a direct link.

Formally, let *z_i_* and *z_j_* denote the final encoder embeddings of the head and tail nodes in the candidate triple (e.g., drug indication-disease). The link prediction head computes:

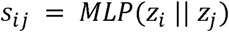

where || denotes the concatenation of the embeddings, and *MLP*(.) consists of four fully connected layers with *ReLU* activation function. The final output is a scalar score *s_ij_* ∈ [0,1], representing the predicted confidence for the link between the head and tail nodes.

## Code availability

All source code and documentation required to reproduce the results reported in this study are publicly available at https://github.com/faren-f/NetMedGPT. An interactive web-based application for candidate drug prioritization for drug repurposing, as well as an interactive chatbot that accepts free-text user queries and returns ranked predictions across all the tasks, is available at https://prototypes.cosy.bio/chatnetmedgpt/. Subnetwork visualizations are currently provided for ten representative diseases; additional disease-specific subnetworks can be generated upon reasonable request where local execution is not feasible.

## Data availability

All datasets used in this study, results, together with the trained model checkpoints, are available at https://cloud.uni-hamburg.de/s/r74Ro8rmQ2sHwsL

## Author contributions

Conceptualization: F.F., J.L., J.B.

Methodology: F.F., J.B.

Investigation: F.F., M.E., D.H., J.B.

Data curation: J.Li, L.L.

Visualization: F.F., S.S.

Software: F.F. (computational framework), S.S. (user interface)

Funding acquisition: J.B.

Project administration: J.B.

Supervision: J.L., J.B.

Writing (Original draft): F.F.

Writing (Review & editing): F.F., S.S., M.E., AM., D.H., J.L., J.B.

We thank J.Li and L.L. for providing curated data and for preliminary analysis of these data.

