## Supplementary information for "NetMedGPT - A network medicine foundation model for extensive disease mechanism mining and drug repurposing"

### Knowledge graph construction

To construct the biomedical knowledge graph (KG) used for training NetMedGPT, we adopted PrimeKG[1], a large-scale heterogeneous KG that integrates curated knowledge from over a dozen biomedical databases. PrimeKG spans 10 node types (Table S1) and 30 relation types (Table S2). The KG is built from a wide range of biomedical sources. These include pharmacological databases such as DrugBank[2] and DrugCentral[3] (which contribute drug-drug, drug-protein, and drug-disease associations, including indications, contraindications, and off-label uses), molecular and genetic resources such as DisGeNET[4], Gene Ontology, and Entrez Gene (covering gene-disease, gene function, and expression patterns), as well as phenotype ontologies like Human Phenotype Ontology[5] and SIDER[6] (providing disease-phenotype and drug-side effect associations). Additional pathway and anatomical context is drawn from sources like Reactome[7], Bgee[8], and UBERON[9], while hierarchical structure among entities (e.g., diseases or functions) is incorporated via ontologies such as MONDO and GO. Details on the individual data sources are available in[1].

### Node feature construction

For each node in the KG, we associated a feature vector, which was used as the initial embedding in NetMedGPT. Of the 10 node types present in the KG, four (drugs, diseases, proteins, and phenotypes) were assigned prior knowledge-based features, while the remaining six node types were randomly initialized with learnable embeddings.

For drugs, diseases, proteins, and phenotypes, we incorporated textual descriptions collected from curated biomedical sources (Table S3). These descriptions were embedded using PubMedBERT, a transformer-based language model pretrained on biomedical literature, to generate semantic representations for each entity. In addition to the textual embedding, drugs were further represented by chemical structure fingerprints. Drugs were first mapped to their SMILES strings via DrugBank identifiers, and circular fingerprints of length 1024 bits were computed using the rcdk package in R. Molecules were parsed using parse.smiles(), and binary fingerprint vectors were generated with get.fingerprint(type = “circular”), where each bit indicates the presence of a specific molecular substructure.

For proteins, we also derived sequence-based embeddings. Entrez Gene identifiers were mapped to UniProt accessions, and full amino acid sequences were retrieved from the UniProt REST API. Each sequence was then embedded using ESM-2 (facebook/esm2_t6_8M_UR50D)[10], a transformer-based protein language model developed by Meta AI.

For the remaining node types, where structured or textual attributes were unavailable, we used randomly initialized embedding vectors. These were treated as trainable parameters and optimized during model training.

### Experimental setup

To evaluate the performance and generalizability of NetMedGPT, we adopted three evaluation strategies inspired by Huang et al.[11], each designed to test the model under increasingly challenging and realistic settings.

**1. Random Link Split**
In this evaluation protocol, edges of each specific type (e.g., drug indication-disease) were randomly partitioned into training (90%), validation (5%), and test (5%) sets, repeating the split five times using different seed numbers {1, 2, 3, 4, 5}. For fair comparison with TxGNN, we used the same splits provided by Huang et al.[11]. While the random link split approach is widely used in the literature, it has important limitations in our context. In the drug repurposing task, for instance, diseases often have multiple therapeutically similar drugs in the KG; hence a random sampling may result in highly similar drug-disease pairs being distributed across training and test sets. Consequently, the model may exploit these similarities, either via shared features or similar connectivity patterns in the KG, making the prediction task easier. Thus, although informative for model benchmarking, this setup may not entirely reflect the real-world challenge of predicting treatments for understudied or rare diseases with few or no known therapies.

**2. Zero-Shot Disease Split**
To better assess generalization, we adopted a zero-shot setting, in which all disease nodes were split into training (90%), validation (5%), and test (5%) groups, ensuring that no drug-disease edges from test diseases were seen during training. This setup simulates drug repurposing for previously unseen diseases. We repeated this evaluation five times with the same splits provided by Huang et al.[11]. However, even in this scenario, test diseases may share similar features, e.g. connectivity or phenotypic profiles, with training diseases, allowing the model to rely on indirect similarities for prediction.

**3. Disease-Area Split**
To further challenge the model's generalizability, particularly for rare diseases that may not share therapeutic or biological connections with others, we employed disease-area split, which is a more stringent strategy. In this setting, we grouped diseases based on their clinical categories (e.g., metabolic, cardiovascular, and cancerous disorders) as defined by Huang et al.[11], and held out entire therapeutic links from disease areas during training. All drug-disease links associated with a given disease area were removed from the training and validation sets and used exclusively for testing. This more realistic and rigorous evaluation tests the model's ability to generalize to entirely unseen clinical domains. As before, we used the same splits provided by Huang et al.[11].

### Negative link generation for evaluation

To evaluate NetMedGPT's ability to distinguish true drug-disease associations from false ones, we constructed a set of negative links during testing. It is important that these negative links are not trivially distinguishable from positive ones, to avoid artificially inflating the model's performance. To address this, we ensured the negative links to have the following conditions.

1. Negative samples were restricted to active tail nodes. For example, in drug repurposing, where the head and tail nodes are a disease and a drug, respectively, the drug nodes should be selected among those that are at least connected to a disease. In particular, in PrimeKG, among 7,957 drugs, only 1,801 are connected to at least one disease via therapeutic indication relationships. The remaining 6,156 drugs have no known therapeutic associations in the KG. Using all drugs for generating negative samples would make the classification task unrealistically easy, as the model could simply learn to discriminate based on drug connectivity. To mitigate this bias, we restricted our negative sampling pool to only those drugs that were already associated with at least one disease in the KG. Such a strategy ensures that all tail nodes in both the positive and negative sets have associated head nodes.

2. The head nodes on negative links were degree-matched with the positive edges. For example, in the drug repurposing task, we matched the number of negative drug samples to the number of positive drug associations for each disease. For a given test disease, we randomly sampled an equal number of drugs (from the therapeutically active pool) that are not known to be associated with that disease. This strategy prevents biases due to node degrees ensuring a fair evaluation setup, where the model must discriminate based on nuanced biological context rather than simple and non-relevant topological features.

### Implementation details

NetMedGPT was implemented in Python (v3.10) using PyTorch (v2.2) as the core deep learning framework. The transformer encoder was built using *torch.nn.TransformerEncoder* and *torch.nn.TransformerEncoderLayer*, with the GELU activation function and multi-head self-attention modules. Optimization was performed using the Adam optimizer, with cross-entropy loss applied over masked tokens during pretraining. For efficient memory management during training, mixed-precision training was enabled using PyTorch's Automatic Mixed Precision (AMP) toolkit, specifically *torch.cuda.amp.autocast* and *GradScaler*. All data loading and batching were handled using *torch.utils.data.DataLoader*.

To compare the performance of NetMedGPT across various biomedical inference tasks, we benchmarked it against recent high-performing GNN-based models that are widely used for heterogeneous KG learning. Specifically, we included three state-of-the-art baselines:

(i) Relational Graph Convolutional Networks (RGCN)[12], which extend standard GCNs to handle multi-relational data by applying relation-specific transformation weights to neighboring nodes. (ii) Heterogeneous Attention Network (HAN)[13], which incorporates both node-level and semantic-level attention mechanisms to learn the relative importance of neighbors and meta-paths in heterogeneous graphs. This allows the model to capture complex semantics in biomedical graphs with multiple node and edge types. (iii) Heterogeneous Graph Transformer (HGT)[14], which employs type-specific attention and message-passing mechanisms to scale

GNNs to large heterogeneous graphs while preserving the relational inductive biases. HGT has demonstrated strong performance in a wide range of knowledge-driven applications, making it a suitable baseline for biomedical KG reasoning. To implement RGCN, HAN and HGT, we used *RGCNConv*, *HANConv* and *HGTConv* layers, respectively, provided in the Pytorch Geometric (vX.X). We trained and evaluated all models using the same training, validation, and test splits as NetMedGPT for fair comparison across five downstream tasks. Hyperparameters for each baseline model were tuned individually using the validation set based on the same optimization method (see next) to ensure optimal performance.

### Hyperparameter optimization

To identify optimal hyperparameters, we used Ray Tune (v2.49.2) with grid search across a predefined search space, evaluating performance on the validation set. The search explored the following hyperparameters: hidden dimension size [100, 200, 300], number of attention heads [5, 10], and number of transformer encoder layers [10, 20, 30]. Each configuration was trained for up to 100 epochs and optimized using the *Asynchronous Successive Halving Algorithm* (ASHA) scheduler, which adaptively allocated computational resources and performed early stopping. Model selection was based on the area under the precision-recall curve (AUPRC) on the validation set. The final model used a hidden dimension of 300, 5 attention heads, and 30 transformer encoder layers, with a batch size of 4000, and a learning rate of $10^{-4}$.

To ensure a fair and robust comparison, we also tuned hyperparameters for all GNN baselines (RGCN, HAN, HGT) using Ray Tune with grid search. The tuning process was repeated across five seed numbers to assess robustness. The search space included: hidden dimension size [10, 20, 30], number of encoder layers [2,3,4], learning rate [0.001, 0.0001], and for HAN and HGT, number of attention heads [2,5]. Each configuration was trained for up to 1000 epochs with early stopping based on performance of AUPRC on validation set, using a patience threshold of five epochs. Training was conducted using the Adam optimizer. Negative sampling and edge splitting strategies were aligned with those used for NetMedGPT to ensure consistency across models.

### Hardware specifications

All experiments were conducted on a high-performance computing node equipped with one NVIDIA H100 NVL GPU (96 GB), two NVIDIA H100 PCIe GPUs (each 81 GB).

**Table S1. Summary of node types in the biomedical knowledge graph (PrimeKG).**

| Node type | Count | Percent in the KG | Source(s) |
|---|---|---|---|
| Drug | 7,957 | 6.15 | DrugBank[15], Drug Central, SIDER |
| Disease | 17,080 | 13.2 | CTD, DisGeNET, Disease Ontology, Drug Central, Human Phenotype Ontology, Mayo Clinic, MONDO Disease Ontology, Orphanet |
| Protein | 27,671 | 21.39 | Bgee, CTD, DisGeNET, DrugBank[15], Entrez Gene, Human Phenotype Ontology, Human PPI Network, Reactome, UMLS |
| Phenotype | 15,311 | 11.83 | DisGeNET, Human Phenotype Ontology, SIDER |
| Biological process | 28,642 | 22.14 | CTD, Entrez Gene, Gene Ontology |
| Cellular component | 4,176 | 3.23 | CTD, Entrez Gene, Gene Ontology |
| Exposure | 818 | 0.63 | CTD |
| Pathway | 2,516 | 1.94 | Reactome |
| Molecular function | 11,169 | 8.63 | CTD, Entrez Gene, Gene Ontology |
| Anatomy | 14,035 | 10.85 | Bgee, UBERON |
| Total number of nodes | 129,375 | 100 | 20 sources |

**Table S2. Summary of edge types in the biomedical knowledge graph (PrimeKG).**

| Edge type | Count | Percent in the KG |
|---|---|---|
| Anatomy-Protein (present) | 3,036,406 | 37.5 |
| Drug-Drug | 2,672,628 | 33.0 |
| Protein-Protein | 642,150 | 7.9 |
| Disease-Phenotype (positive) | 300,634 | 3.7 |
| Biological process-Protein | 289,610 | 3.6 |
| Cellular component-Protein | 166,804 | 2.1 |
| Disease-Protein | 160,822 | 2.0 |
| Molecular function-Protein | 139,060 | 1.7 |
| Drug-Effect | 129,568 | 1.6 |
| Biological process-Biological process | 105,772 | 1.3 |
| Pathway-Protein | 85,292 | 1.1 |
| Disease-Disease | 64,388 | 0.8 |
| Drug-Disaese (contraindication) | 61,350 | 0.8 |
| Drug-Protein | 51,306 | 0.6 |
| Anatomy-Protein (absent) | 39,774 | 0.5 |
| Phenotype-Phenotype | 37,472 | 0.5 |
| Anatomy-Anatomy | 28,064 | 0.3 |
| Molecular function-Molecular function | 27,148 | 0.3 |
| Drug-Disease (indication) | 18,776 | 0.2 |
| Cellular component-Cellular component | 9,690 | 0.1 |
| Phenotype-Protein | 6,660 | 0.1 |
| Drug-Disease (off-label use) | 5,136 | 0.1 |
| Pathway-Pathway | 5,070 | 0.1 |
| Exposure-Disease | 4,608 | 0.1 |
| Exposure-Exposure | 4,140 | 0.1 |
| Exposure-Biological process | 3,250 | 0.04 |

| | | |
|---|---|---|
| Exposure-Protein | 2,424 | 0.03 |
| Disease-Phenotype (negative) | 2,386 | 0.03 |
| Exposure-Molecular function | 90 | 5e-04 |
| Exposure-Cellular component | 20 | 25e-05 |
| Total number of edges | 4,050,249 | 100.0 |

**Table S3. Node descriptions and their sources.**

This table summarizes the types of node-level textual descriptions incorporated into NetMedGPT as input features. For each node type, the table lists the specific content used along with the corresponding data sources.

| Node types | Description | Source(s) |
|---|---|---|
| Drug | SMILES, drug description containing description, mechanism of action. | DrugBank, PrimeKG (Chandak, Huang, and Zitnik, 2023) |
| Disease | Disease descriptions containing mondo name, mondo definition, UMLS description, orphanet definition, orphanet clinical description | PrimeKG (Chandak, Huang, and Zitnik, 2023), |
| Protein | Protein structure, protein description | UniProt, NedRex[16] |
| Phenotype | Phenotype description | NedRex[16] |

**Table S4. Impact of node attributes on model performance.**

To assess the contribution of node-level features, we compared the performance of NetMedGPT with and without node features. Including node features led to consistent improvements across all evaluation metrics.

| Relations | NetMedGPT w/o features (AUPRC) | NetMedGPT (AUPRC) |
|---|---|---|
| Indication | $0.95 \pm 0.00$ | $0.97 \pm 0.00$ |
| Contraindication | $0.91 \pm 0.00$ | $0.93 \pm 0.00$ |
| Off label use | $0.95 \pm 0.00$ | $0.97 \pm 0.00$ |
| Drug-Protein | $0.90 \pm 0.00$ | $0.95 \pm 0.00$ |
| Drug-Effect | $0.92 \pm 0.00$ | $0.93 \pm 0.00$ |

**Table S5. Number of disease nodes with indication and contraindication edges in each disease area.**

| **Disease area** | **Number of diseases with indication link** | **Number of diseases with contraindication link** |
|---|---|---|
| Diseases of cell proliferation | 152 | 85 |
| Mental health diseases | 29 | 36 |
| Cardiovascular diseases | 41 | 80 |
| Diseases of anemia | 8 | 12 |
| Adrenal gland diseases | 6 | 4 |
| Autoimmune diseases | 13 | 11 |
| Metabolic disorders | 28 | 23 |
| Diabetes | 2 | 3 |
| Neurodegenerative diseases | 12 | 12 |

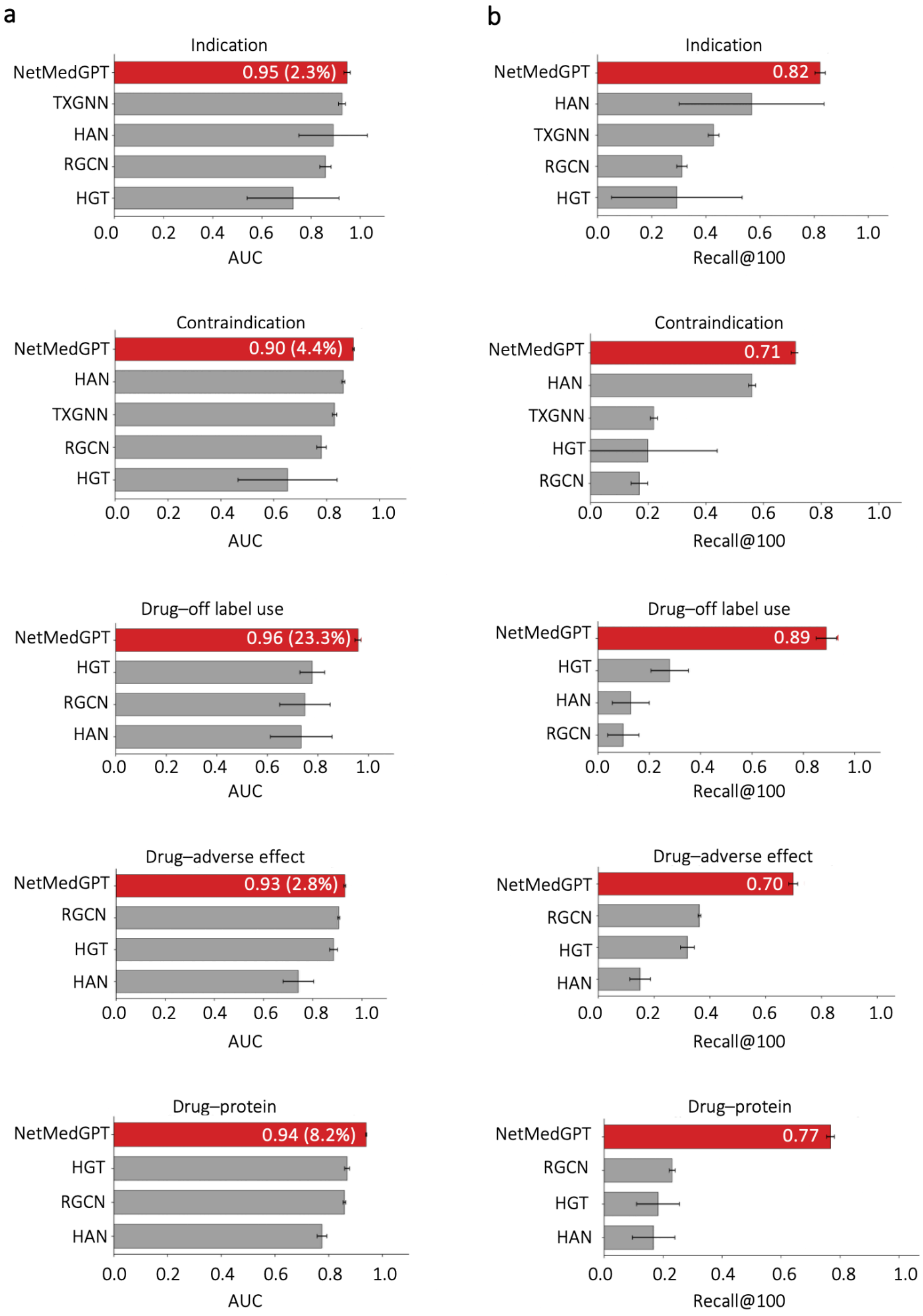

**Figure S1. AUC and Recall@100 performance across biomedical prediction tasks.**
Comparison of NetMedGPT with baseline models on the prediction of drug indication, contraindication, off-label use, adverse drug reaction, and drug target interaction. Bars represent AUC (left) and Recall@100 (right) for each method. NetMedGPT consistently achieves superior performance in both AUC and Recall@100 across all evaluated tasks.

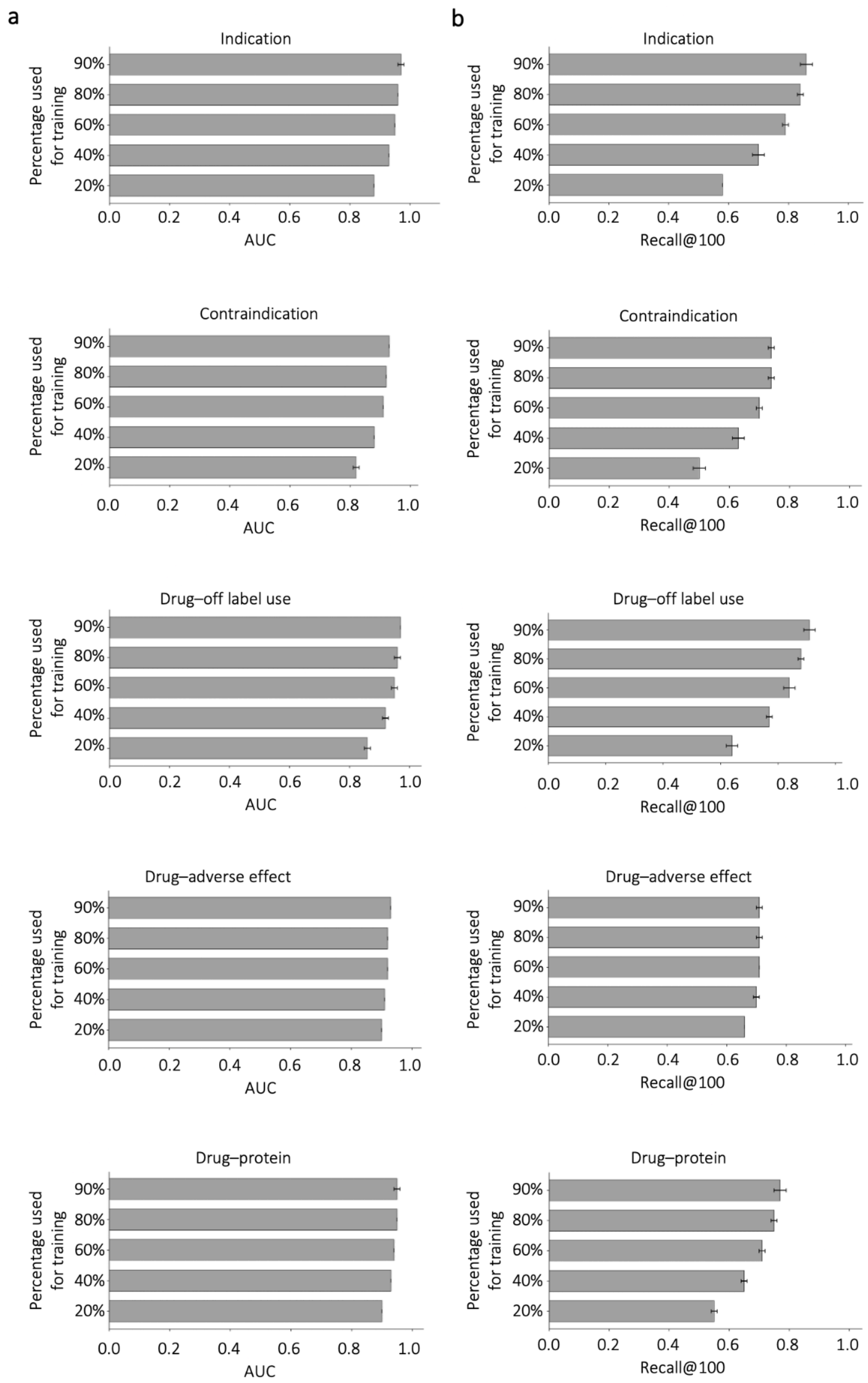

**Figure S2. Model performance across different training data proportions.**
Performance of the model evaluated on five prediction tasks, indication, contraindication, off-label use, adverse effect, and drug-protein, using varying percentages of training data (20-90%). (a) Area under the receiver operating characteristic curve (AUC); (b) Recall@100. Bars represent mean performance across runs, and error bars indicate standard deviation.

Model performance generally improves with increasing training data, demonstrating robustness and scalability across multiple biomedical prediction tasks.

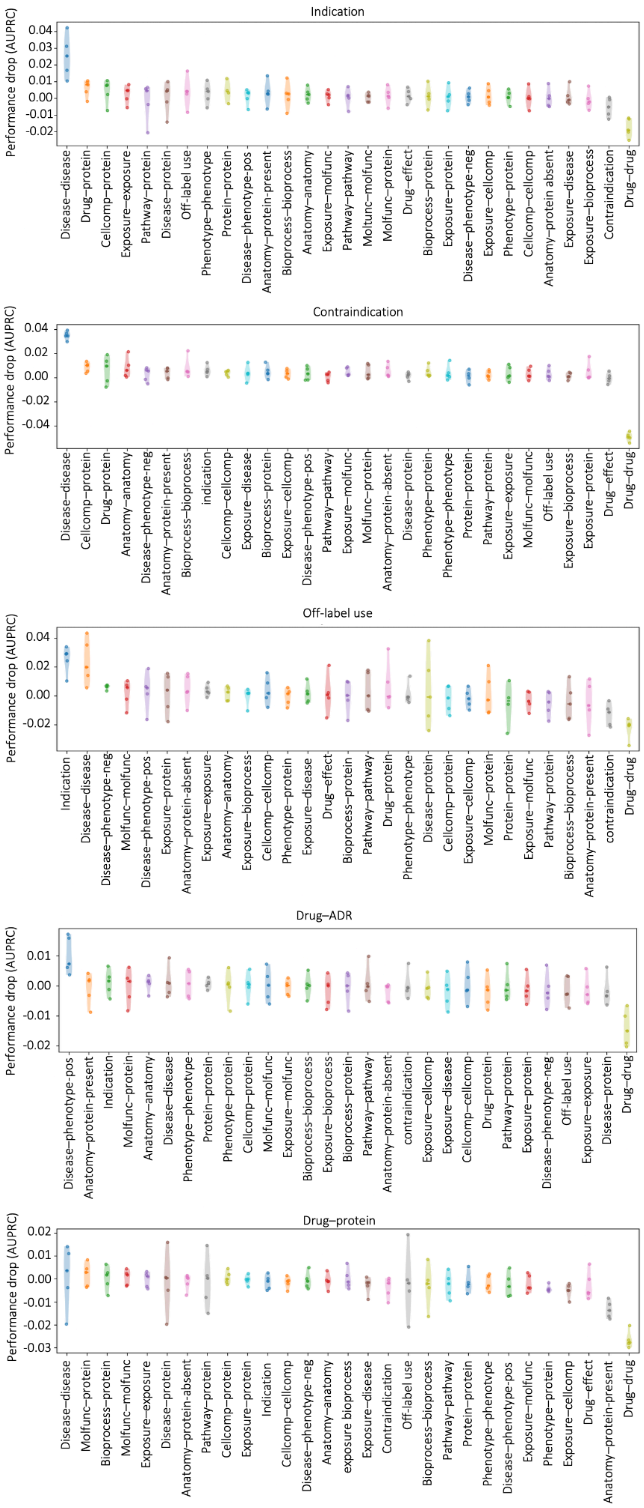

**Figure S3. Robustness analysis of NetMedGPT following edge-type ablation.**
The impact of removing individual relation types from the knowledge graph on model performance across different prediction tasks. Each point represents the performance drop ($\Delta AUPRC$) observed when excluding a specific relation type during training. The results show that the overall performance of NetMedGPT is stable against edge-type removal.

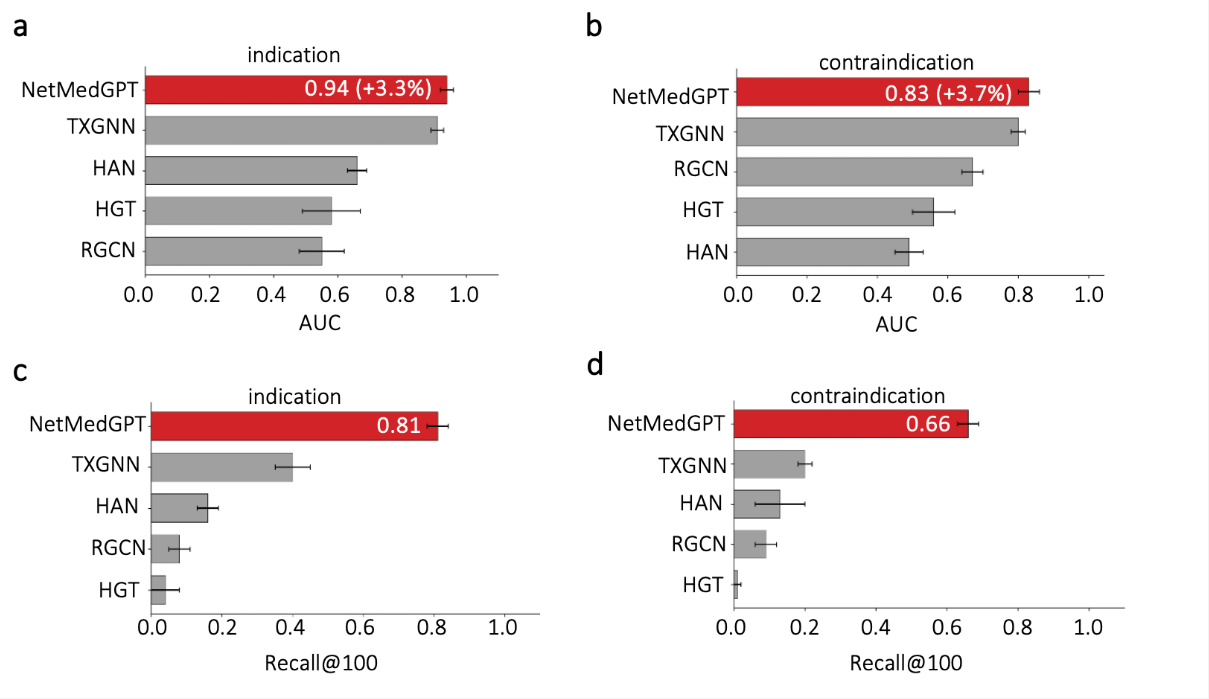

**Figure S4. AUC and Recall@100 performance across indication and contraindication tasks under the zero-shot split.**
Bars represent AUC (left) and Recall@100 (right) for each method. NetMedGPT consistently outperforms all baseline models across both tasks and evaluation criteria, demonstrating superior generalization under the zero-shot setting.

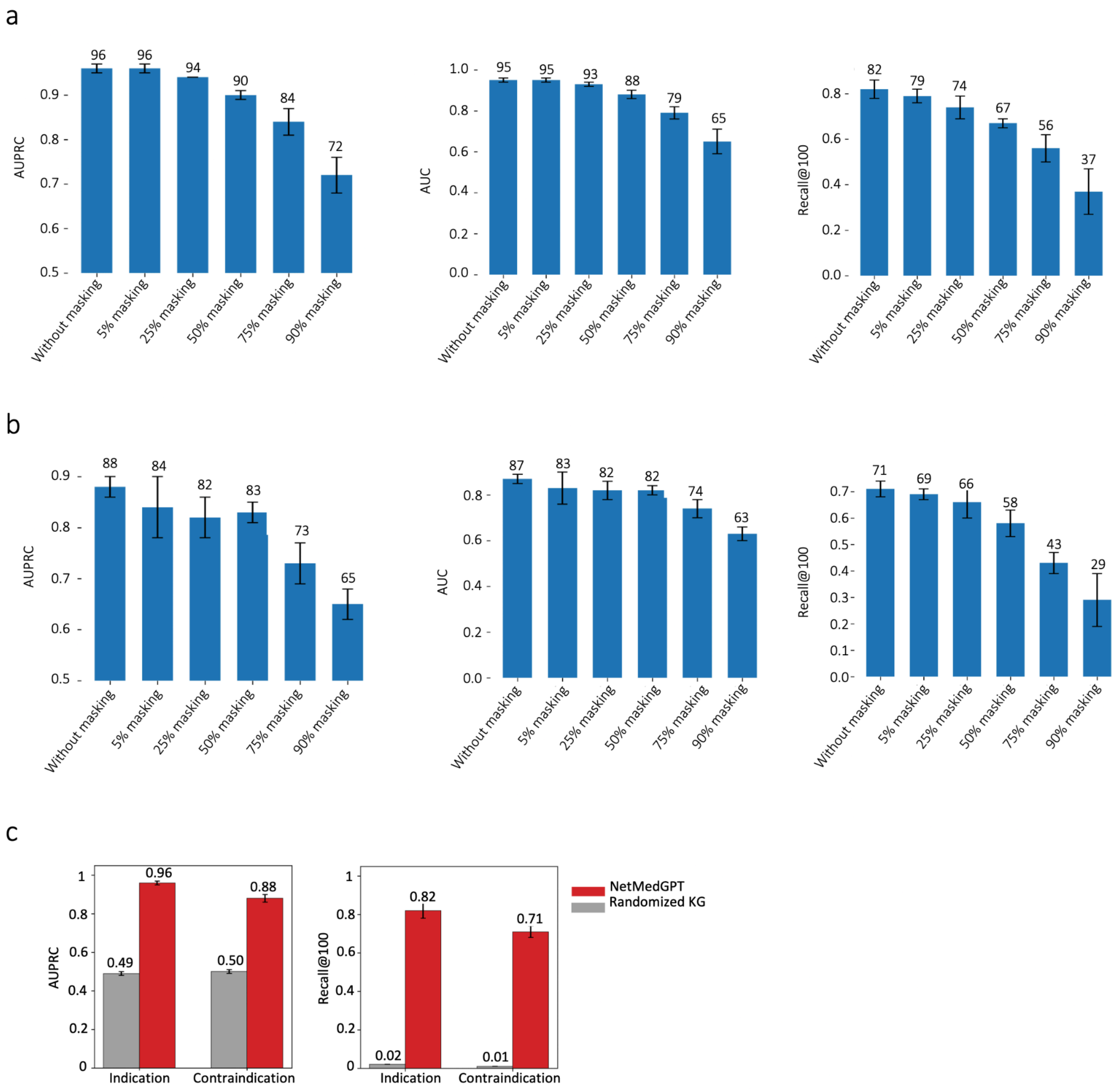

**FigureS5. Performance stability of NetMedGPT.**

The performance of NetMedGPT after randomly removing 5%, 25%, 50%, 75%, and 90% of the edges in the KG for a) indication and b) contraindication tasks. c) The performance of NetMedGPT trained on PrimeKG and randomized versions of PrimeKG.

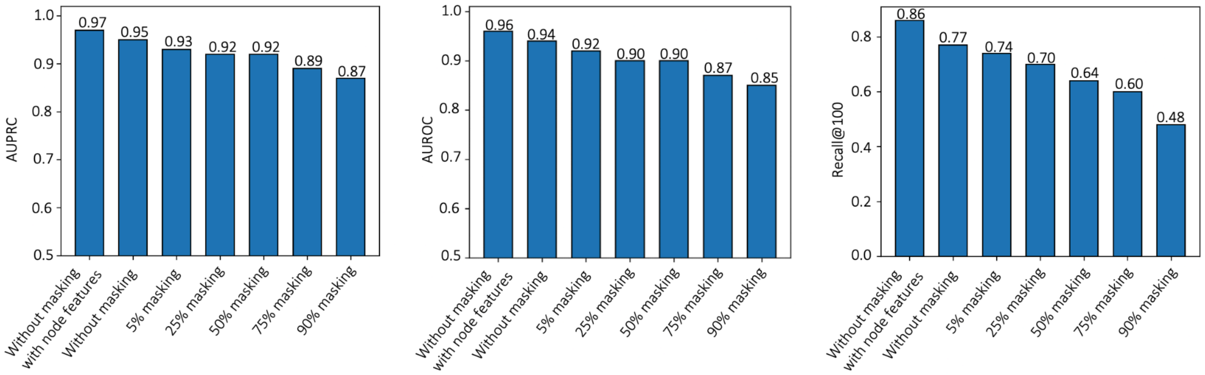

**Figure S6. Performance stability of NetMedGPT under masking of biological neighbors.**
We evaluated the sensitivity of NetMedGPT to the removal of 1-hop biological neighbors of diseases. Specifically, we measured performance with increasing levels of masking, removing 5%, 25%, 50%, 75%, and 90% of the 1-hop biological context. While performance showed a gradual decrease, as expected, it remained stable, demonstrating the model's robustness.

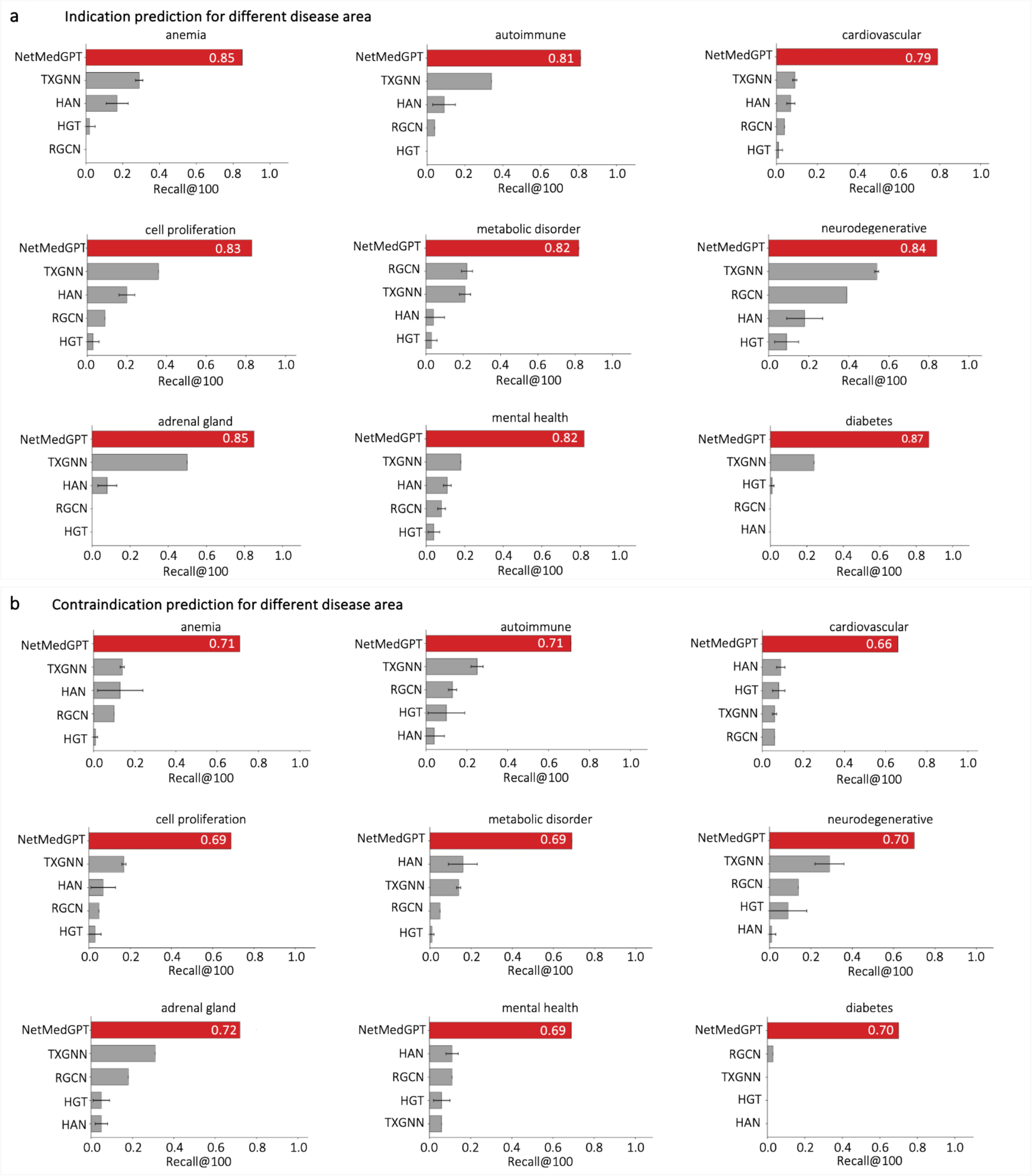

**Figure S7. Recall@100 performance across indication and contraindication prediction tasks under the disease-area split.**

Bars represent Recall@100 for each method. NetMedGPT consistently outperforms baseline models across all disease areas for both tasks.

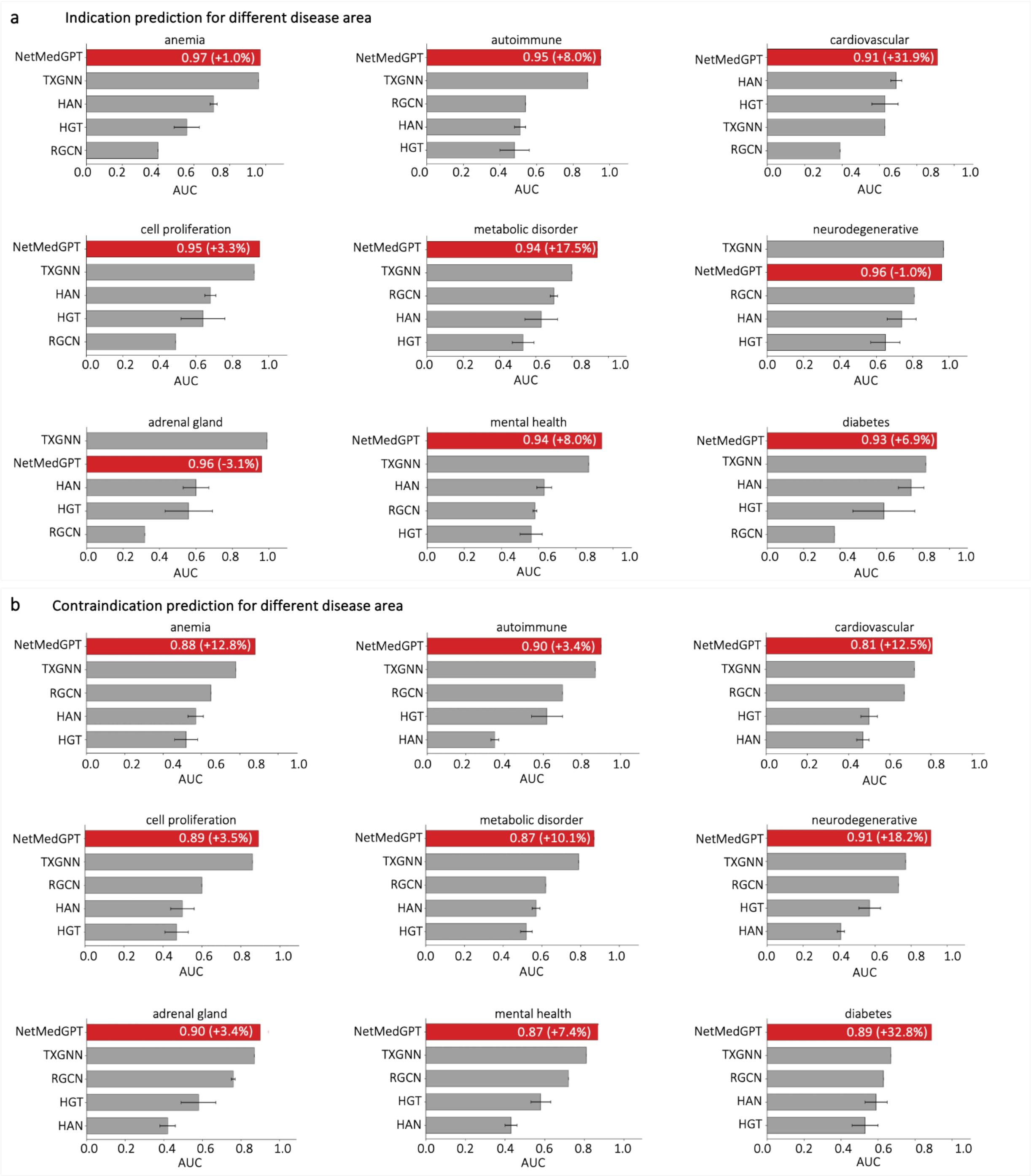

**Figure S8. AUC performance across indication and contraindication prediction tasks under the disease-area split.**

Bars represent AUC for each method. NetMedGPT consistently outperforms baseline models a) across 7 out of 9 disease areas for indication prediction and b) all disease areas for contraindication prediction.

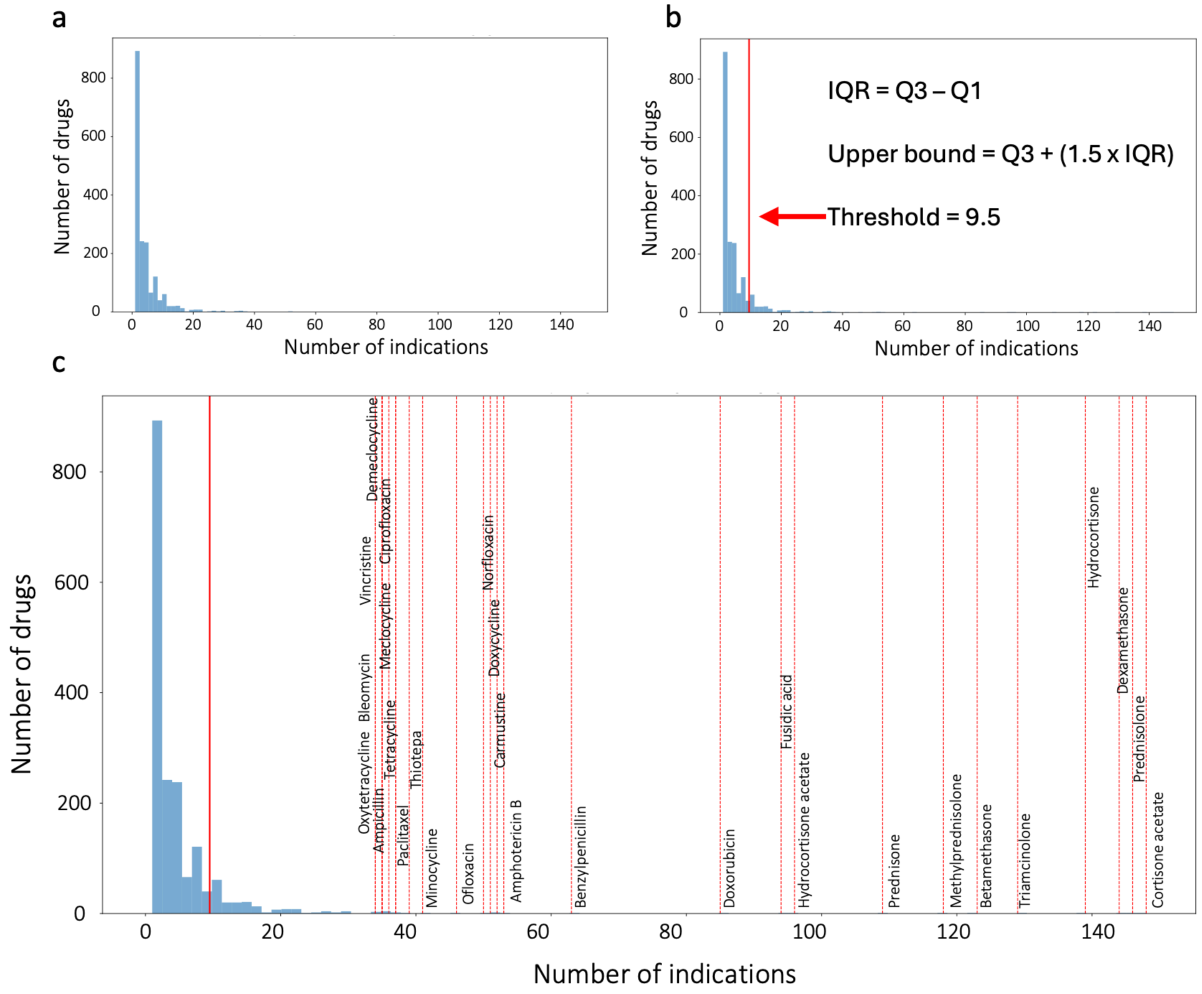

**Figure S9. Identification and removal of indication-count outliers.**

(a) Distribution of the number of indications per drug, showing a highly right-skewed distribution.

(b) Determination of the outlier threshold using the Tukey method. The upper bound was defined as Q3 + 1.5 × IQR, yielding a threshold of 9.5 indications (red line).

(c) Some drugs exceeding this threshold are shown along the x-axis. These drugs primarily belong to broad therapeutic classes, glucocorticoids, antibiotics, antifungals, and cytotoxic chemotherapies, that act through non-specific or class-wide mechanisms unrelated to the disease-focused biological pathways. Because they form clear statistical and mechanistic outliers relative to the rest of the drugs, they were excluded from downstream analyses.

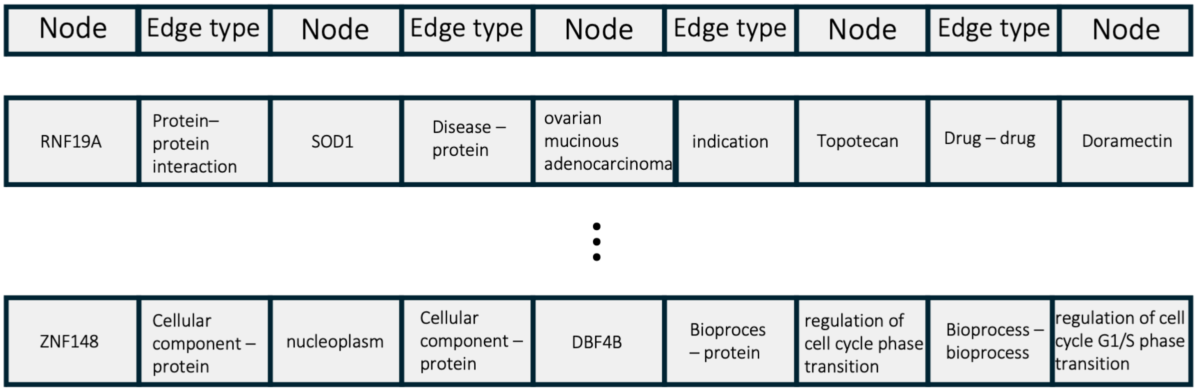

**Figure S10. Pseudo-sentence construction from random walks over the knowledge graph.** Each random walk begins at a node and proceeds through the heterogeneous knowledge graph based on edge connectivity until a predefined length is reached. For every consecutive pair of nodes in the walk, the connecting edge type is inserted to form a structured pseudo-sentence. These pseudo-sentences serve as input sequences for NetMedGPT's masked token prediction task.
